# HCF1 orchestrates O-GlcNAcylation and affinity-dependent transcription through extended molecular determinants and register-shifted binding

**DOI:** 10.64898/2026.03.19.712392

**Authors:** Mihkel Örd, Sydney A. Porto, Amelia Barclay, Mingxuan Jiang, Liz Kogan, Michelle Leiser, Pau Creixell

## Abstract

We recently identified Host Cell Factor 1 (HCF1), a transcriptional co-regulator discovered more than thirty years ago, as a cancer dependency. To further understand its molecular functions and expand its known interactome, here, we screened a proteome-wide library of candidate HCF1 binding peptides and identified previously uncharacterized canonical and non-canonical HCF1-binding partners. Through deep mutational scanning (DMS) screening, we uncovered an extended set of molecular determinants of binding and show how mutations outside its previously established interacting residues impact binding affinity. Next, we uncover non-canonical HCF1 binders with an extended register-shifted two-amino acid sequence between their “anchor” histidine and tyrosine amino-acid residues, which we show critically contributes, in an affinity-dependent manner, to the downstream transcriptional activity of IRF1. Our data also shows that HCF1 promotes O-GlcNAcylation of the majority of its transcriptional binders. Overall, our results significantly expand the number and diversity of HCF1 binders and propose an enhanced mechanistic understanding of how HCF1 orchestrates transcription and O-GlcNAcylation.

## Introduction

Host Cell Factor 1 (HCF1) was discovered in 1993, as its name suggests, as an accessory protein in host cells that, upon viral infection, becomes hijacked to promote the transcription of viral genes^1^. Through its Kelch domain, HCF1 interacts with a number of proteins historically defined as containing a consensus D/EHxY^2–4^ sequence. This HCF1 binding consensus sequence has been found in over 30 human proteins, with candidate HCF1-binding sequences identified in many more^3^. Through these interactions, HCF1 promotes the transcriptional activation of genes regulating cell cycle^5,6^, metabolism^7^, stress responses^8^, and embryonic stem cell pluripotency^9,10^ (**Fig. 1A, S1A**).

**Figure 1.**
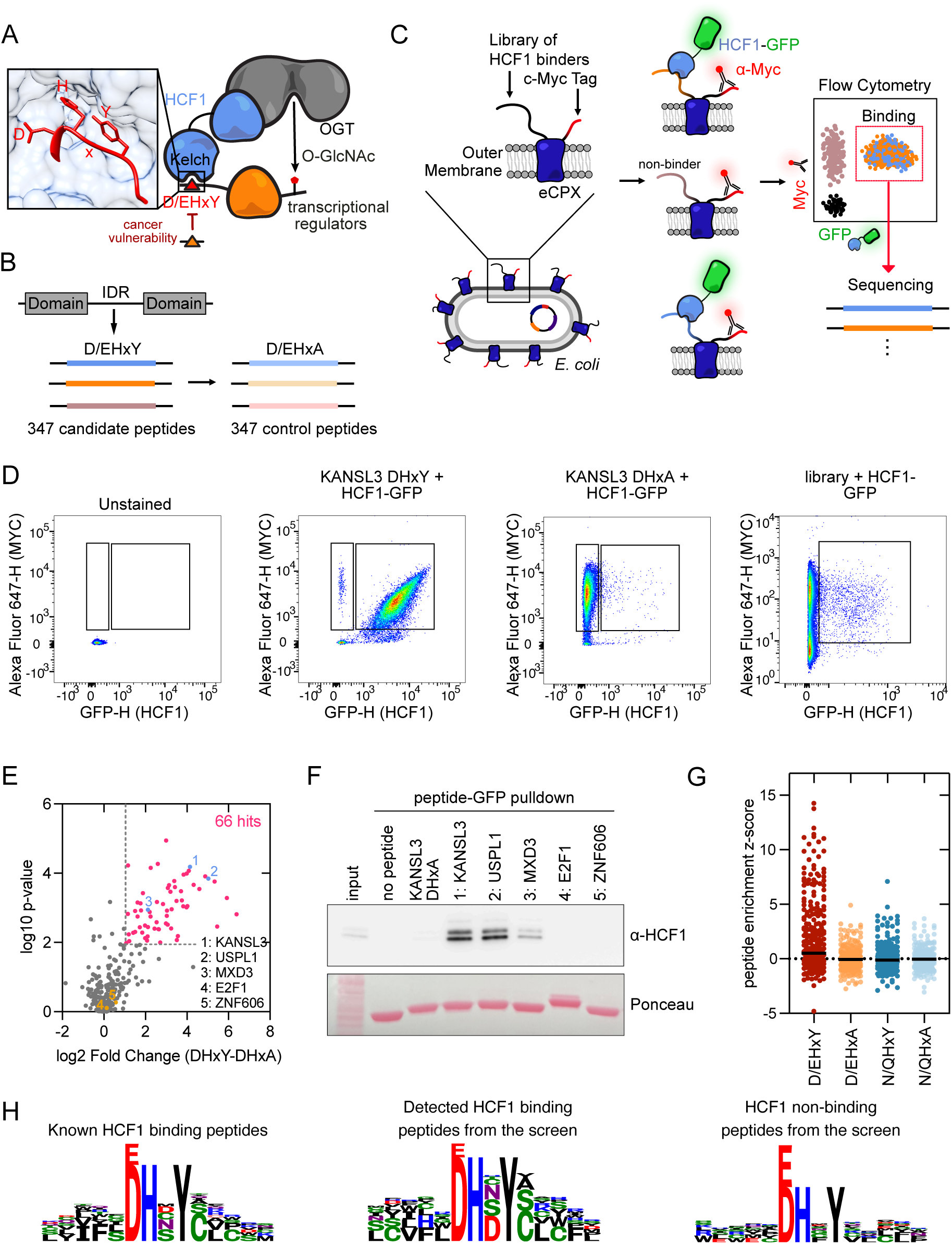
HCF1 binds a subset of D/EHxY peptides. (**A**) Schematic model of HCF1-OGT complex binding to D/EHxY peptides. An AlphaFold3 model of KANSL3 DHSY peptide bound to HCF1 is shown. (**B**) Design of proteome-wide HBM library by assembly of D/EHxY-containing peptides from the intrinsically disordered regions (IDRs) of human proteins. (**C**) Scheme of bacterial surface peptide display experiment used to measure binding of Kelch domain of HCF1 EGFP fusion to *E. coli* cells displaying HCF1 binding peptides fused to MYC-tagged eCPX protein. (**D**) Flow cytometry plots showing HCF1(Kelch)-EGFP and Alexa-647 anti-MYC staining of peptide displaying *E. coli* cells. A representative experiment of two biological replicates is shown. (**E**) Corrected log2 fold change and p-values of D/EHxY peptides for HCF1(Kelch)-EGFP binding in pooled peptide display experiments compared to their alanine control peptides. (**F**) Western blot of a peptide pulldown experiment testing the indicated peptide GFP fusions in pulling down HCF1 from HCT116 cell lysate. A representative example of two biological replicates is shown. (**G**) Distribution of enrichment scores for HCF1(Kelch)-EGFP binding in a pooled peptide display experiment using libraries of 347 D/EHxY peptides or 259 N/QHxY peptides together with their D/E/N/QHxA negative control mutants. (**H**) Sequence logos showing the positional preferences in different groups of D/EHxY peptides.

In a recent proteome-wide screen for cancer dependencies on specific protein-protein interactions, we identified the Kelch pocket of HCF1 as a cancer dependency and potential therapeutic target because, despite being a pan-essential gene, different cancer cell lines showed significant variability in their sensitivity to HCF1 pocket inhibition (**Fig. 1A**)^11^. In addition to this therapeutic window, others have shown that the HCF1-binding sequence in MYC is required for its oncogenic activity which has, in turn, increased interest in targeting the HCF1-MYC interaction as a therapeutic opportunity in MYC-driven cancers^12^. Thus, given its role not only in viral infection, but also in other diseases such as cancer, where it is often upregulated and has been linked to metastatic spread^13–15^, it is becoming important to understand the interaction partners and molecular function of HCF1.

Despite this significant and increasing interest and the number of proteins that have been identified as binders of HCF1^3,16^, the lack of structural and systematic identification and characterisation of its binders limits our understanding of its function. While the early alanine-scanning experiments identified a consensus D/EHxY sequence^2^, subsequent studies have found additional targets (including BAP1, MYC1, and NRF1)^8,12,17^ with a divergent N/QHxY HCF1-binding sequence. Thus, we hypothesized that the full diversity of HCF1 binding proteins, the complete set of molecular determinants of binding and interaction affinity, and how binding and affinity contribute to the signalling output of HCF1 and its interaction partners remain unknown and yet important biological questions to solve.

Indeed, even for one of its best-known functions as transcriptional co-regulator, the molecular mechanism by which HCF1 regulates transcription remains poorly understood. While HCF1 is not believed to directly bind DNA or possess direct catalytic activity, it has been suggested that HCF1 may act as a scaffold protein bringing together DNA-binding proteins and proteins involved in post-translational modifications^18^. As a primary example, HCF1 is known to form a stable complex with O-linked N-acetylglucosamine (O-GlcNAc) transferase, OGT, driving its function to such an extent that over 50% of nuclear OGT is believed to be bound to HCF1^19^. The fact that OGT interacts with the C-terminal half of HCF1^19^ may be mechanistically relevant as it leaves the Kelch domain of HCF1 available to interact with other proteins thereby forming distinct OGT complexes (**Fig. 1A**). Moreover, O-GlcNAc-modified proteins appear to be enriched at the promoters of actively transcribed genes and this modification is believed to be a positive regulator of transcription^20^. All this, together with the fact that HCF1 promotes the O-GlcNAcylation of NRF1 and PGC1ɑ by OGT^7,8^, that HCF1 stabilizes OGT, and that the loss of HCF1 has been associated with decrease in O-GlcNAc-modified proteins^21^, made us hypothesize that HCF1 may function as a broad regulator of O-GlcNAcylation.

Given the key role of the Kelch domain pocket in HCF1 function, here, we systematically studied and molecularly characterized its interactions by screening a proteome-wide library of candidate canonical and non-canonical HCF1-binding partners. Through these screens, we find that only a subset of consensus-containing peptides are capable of productive HCF1 binding, which further motivated us to map additional molecular determinants of binding by deep mutational scanning (DMS). Through our subsequent experiments, we uncover previously unrecognized molecular determinants of binding affinity and non-canonical binders with an extended register-shifted intermediate binding sequence. Capitalizing on this new molecular understanding, we quantitatively assess the impact of differential binding on the transcriptional response downstream of HCF1. To decipher the degree to which its function is driven by its association with OGT, we finally test the role of HCF1 in mediating O-GlcNAcylation of its binding partners. Our study, thus, provides not only a comprehensive resource systematically cataloguing canonical and non-canonical HCF1-binding proteins, but also a characterization of the rules determining their binding strengths and downstream effects.

## Results

### Our high-throughput proteome-wide peptide display assay enables the systematic study of HCF1 binders and reveals that only a subset of consensus peptides are functional

To study HCF1 binding motifs in a high-throughput manner, we set up bacterial surface peptide display experiments, where the peptides are expressed as an eCPX-MYC fusion protein, resulting in their display on the surface of *E. coli* cells^22^ (**Fig. 1B-C**). We then mix these cells with an Alexa-647-labelled anti-MYC antibody to confirm successful display of the peptide-eCPX-MYC and purified the Kelch domain of HCF1 tagged with EGFP, HCF1^Kelch^ ^domain^-EGFP, to detect HCF1 binding. The observation that HCF1^Kelch^ ^domain^-EGFP stains *E. coli* cells displaying the DHxY peptide from KANSL3, but not *E. coli* cells displaying a non-binding DHxA control peptide (**Fig. 1D**) suggests that our assay can effectively discriminate binders from non-binders.

Given that the known HCF1-binding sequences reside in intrinsically disordered regions (IDRs) of proteins and, in most cases, match a consensus D/EHxY sequence^3^, we constructed a proteome-wide library containing all 347 peptides within the human proteome containing this sequence in IDRs (**Fig. 1B**). Next, we displayed this proteome-wide library (alongside their counterpart 347 D/EHxA negative control peptides) in the exposed N-terminal region the eCPX scaffold, incubated these bacterial cells with purified HCF1^Kelch^ ^domain^-EGFP and used FACS to sort for EGFP-positive cells. The finding that the majority of *E. coli* cells in this library did not bind HCF1^Kelch^ ^domain^-EGFP suggests that a large fraction of D/EHxY-containing peptides in the proteome are not *bona fide* HCF1 binders, or that if they bind, they do so with a weaker affinity that is below the limit of detection of our assay (**Fig. 1D**). Finally, to identify the HCF1-binding peptides within our proteome-wide library, we performed deep sequencing from our GFP-positive sorted cells (alongside our input library) and obtained enrichment scores that, as has been shown previously^23^, are proportional to their target binding capacity (**Fig. S1B**).

Comparing our wild-type peptides to their DHxA negative control peptides, we find that 66 of our 347 proteome-wide library peptides bind to HCF1 (**Fig. 1E**). Out of these 66 *bona fide* HCF1 binders, we found that, in addition to the 25 previously identified HCF1 binders, many of the 41 previously uncharacterized HCF1-binding peptides are transcriptional regulators, including DNA-binding, chromatin-modifying, and SUMOylation-related proteins (**Table S1, Fig. S1C**). To further investigate why our assay had not detected binding to 11 known HCF1-binding proteins, we performed deep sequencing from our initial library-expressing MYC-positive population, which revealed that a small number of peptides (including the HCF1-binding peptides from PHF8 and THAP1) display poorly (**Fig. S1D**). Thus, we concluded that for rare cases where specific peptide sequences result in poor display this would prevent us from effectively detecting these peptides as enriched in our assay. To further assess the specificity of our assay, we compared our peptide display data with an HCF1 co-IP dataset that we had previously generated^11^ and found that our enrichment scores for HCF1-binding proteins were significantly higher than the enrichment scores for non-binding D/EHxY-containing proteins (based on our co-IP experiments) (**Fig. S1E**). This strongly suggests that our false positive rate is low and, as a consequence, the specificity of our pooled display assay is high. To orthogonally validate our candidate HCF1 binders, we performed peptide pulldown experiments using five peptides with varying enrichment scores (**Fig. 1E**). These experiments not only validated the previously uncharacterized candidates USPL1 and MXD3 as *bona fide* HCF1 binders but also supported that they bind HCF1 with a comparable affinity as the known HCF1 binder KANSL3. Moreover, the fact that we did not detect any pulldown signal for the two peptides with low enrichment scores, namely E2F1 and ZNF606, (**Fig. 1F**) further supports that our high-throughput peptide display assay also presents a high degree of specificity.

In an attempt to more broadly cover the whole known HCF1-binding repertoire, we screened a second proteome-wide library for peptides containing the second consensus N/QHxY sequence. The fact that we obtained fewer enriched binders from this proteome-wide library than from the initial D/EHxY library, suggests that N/QHxY peptides tend to either bind with weaker affinity or not bind at all (**Fig. 1G, Fig. S1F**). Despite this observation, the fact that some of the established N/QHxY binders, BAP1 and members of the MYC family^12,17^, are among top enrichment scorers in our screen, suggests that our assay is also capable of recovering and identifying *bona fide* N/QHxY binders (**Fig. S1G**).

Taken together, these experiments propose a model in which only a subset of peptides with the D/E/N/QHxY peptide sequences result in productive HCF1 binding. Moreover, the sequence logos we obtained from our HCF1-binding peptides (also partially consistent with logos obtained from known HCF1 binders) suggest that there are additional molecular determinants of binding, selectivity in both N- and C-terminal flanking residues, and limited tolerated residues in the intermediate “x” position between the “anchor” histidine and tyrosine residues of HCF1-binding partners (**Fig. 1H, S1A**).

### Deep mutational scanning (DMS) identifies extended molecular determinants of binding in the intermediate, N- and C-terminal flanking residues of HCF1 binding partners

Given the lack of structural information around the interaction between the Kelch domain of HCF1 and its interacting peptides^24^, we used AlphaFold3 to computationally model HCF1 binding and guide the design of additional individual and libraries of peptides that would help us further elucidate key molecular mechanisms driving HCF1 binding. In these models, the “anchor” residues part of the consensus D/EHxY sequence are buried when bound to HCF1 as expected (**Fig. 2A-B, S2A**). However, in addition to these “anchor” residues, several additional N-and C-terminal residues appear to be making significant contacts, and be buried, when bound to HCF1 in many of these AlphaFold3 complexes (**Fig. 2A-B, S2A**). These computational models, thus, suggest that additional interactions may drive HCF1 binding.

**Figure 2.**
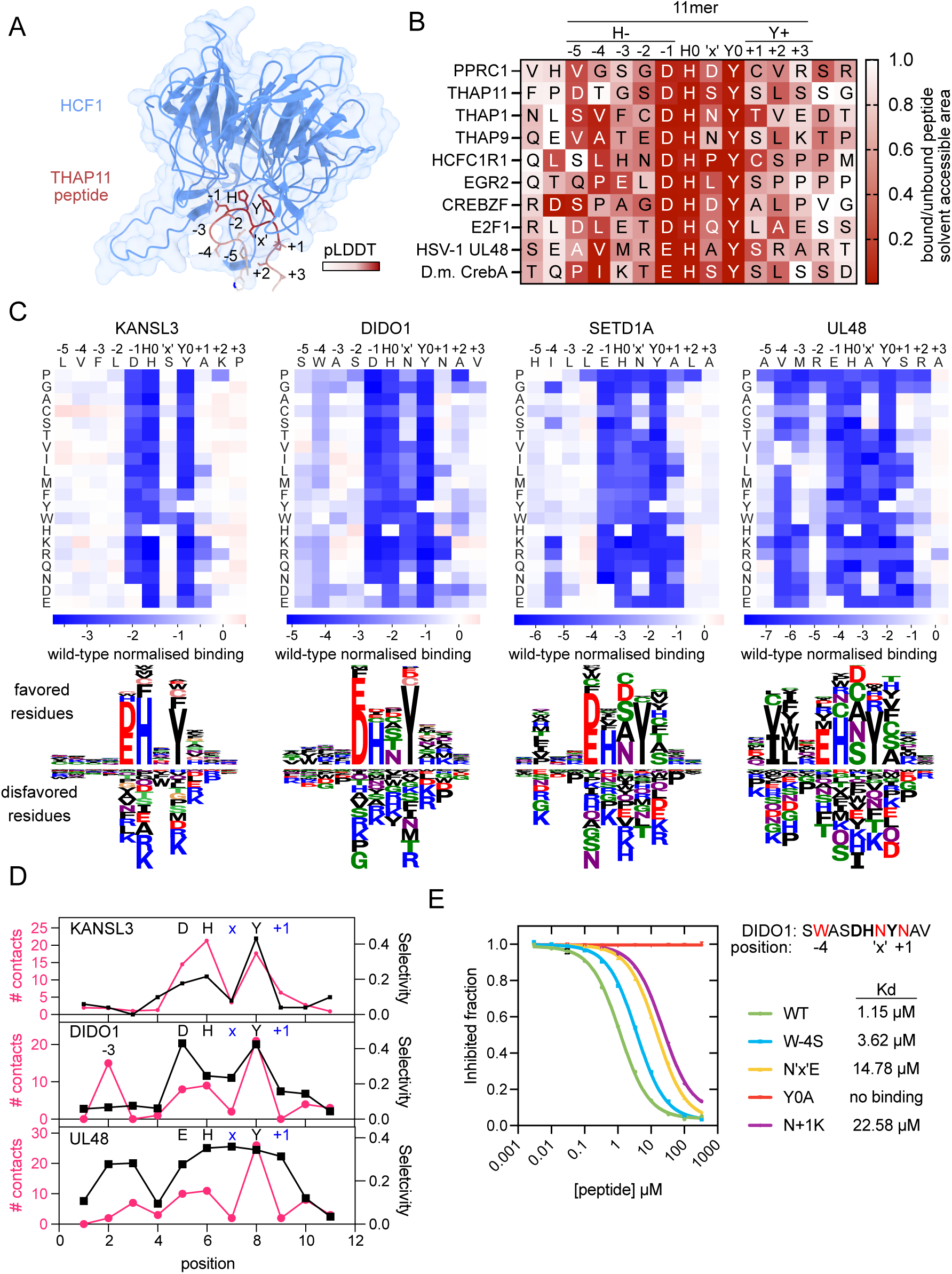
Deep mutational scanning reveals the molecular determinants of binding of HCF1 binding motifs. (**A**) AlphaFold3 model of HCF1 Kelch domain (blue) and THAP11 DHxY peptide complex. The HCF1 153-169 β-sheets are not shown in order to visualize the peptide. (**B**) Alignment of HCF1 binding peptides with the relative solvent accessible areas (peptide in complex with HCF1 compared to unbound peptide) shown for each residue. (**C**) Wild-type-corrected HCF1 binding scores measured in bacterial peptide display with deep mutational scanning libraries of KANSL3, DIDO1, SETD1A, and UL48 peptides. Below, sequence logos showing favored and disfavored residues for HCF1 binding in each position of the peptides. (**D**) The Gini index (selectivity) showing inequality between residues in a position, calculated from DMS data, and the number of van der Waals contacts of the residue with HCF1 in AlphaFold3 model. (**E**) Dose-response curves showing the ability of different DIDO1 peptide variants as competitor peptides to displace a FITC-labeled HCF1 binding peptide from HCF1, measured by fluorescence polarization (n = 2 biological replicates).

To test this hypothesis, we designed and screened 11-mer peptide libraries by deep mutational scanning (DMS). To facilitate our analysis and its interpretation, we numbered N-terminal peptide residues as negative residues relative to the “anchor” histidine residue (e.g., H-1) and C-terminal peptide residues as positive residues relative to the “anchor” tyrosine residue (e.g., Y+1), while labeling the residue between them as the intermediate “x” residue (**Fig. 2B**). Following this scheme, we created DMS peptide libraries containing all possible 20 amino acids in the 11 residues between H-5 to Y+3 using four different HCF1-binding peptides as sequence backgrounds, namely KANSL3, DIDO1, SETD1A, and UL48. By choosing these peptides as background, we also aimed to cover diversity within the H-1 acidic residue position, where KANSL3 and DIDO1 encode an aspartate (D) residue, while SETD1A and UL48 encode a glutamate (E) residue. As earlier, we screened these libraries for HCF1 binding using bacterial peptide display and performed deep sequencing to obtain enrichment scores for all peptide variants in our library (**Fig. S2B-C**). Across all starting backgrounds, in addition to non-binding variants, we identified peptide variants yielding both improved and weakened HCF1 binding compared to their wild-type counterparts (**Fig. 2C**). To rule out the possibility that our observed enrichment was driven by varying display efficiency in our peptide variants (instead of true HCF1 binding), we sorted and performed deep sequencing on the MYC-positive population from our UL48 DMS library-expressing bacterial culture (**Fig. S2D**). The fact that the enrichment scores obtained from HCF1 pulldown did not correlate with display (**Fig. S2E**) suggests that our enrichment scores are driven by HCF1 binding.

While regardless of peptide background, we observed the strongest selectivity in the “anchor” acidic, histidine and tyrosine positions, where no other residues were tolerated, several additional positions also showed clear signs of specificity and binding constraints (**Fig. 2C**). These results are also consistent with our earlier observation that a large fraction of D/EHxY proteome-wide library peptides showed no evidence of HCF1 binding (**Fig. 1E**). Different starting peptides, however, showed different tolerance to mutations, with more variation allowed in KANSL3 than in DIDO1, SETD1A, or UL48 (**Fig. 2C**). Conversely, we also identified common features, including the fact that (even in the more mutation-tolerant background of the KANSL3 peptide) we found proline to be strongly disfavoured in the Y+2 position, presumably because its cyclic side chain makes it incompatible with the peptide backbone torsion required for productive HCF1 binding. In all peptide backgrounds, we also observed strong selectivity in the intermediate ‘x’ and C-terminal Y+1 residues. In the DIDO1, SETD1A, and UL48 backgrounds, only small residues were tolerated at the intermediate ‘x’ position and, in the more mutation-tolerant background of the KANSL3 peptide which encodes relatively low selectivity at this position, we similarly saw large, aromatic residues being disfavoured. In the C-terminal Y+1 position, all peptides disfavor charged residues, whereas in some contexts, glutamine (Q), leucine (L), and tryptophan (W) are also disfavored. While the degree of selectivity varied in the background of each peptide, for DIDO1, SETD1A, and UL48 we also observed significant specificity in the N-terminus of the binding peptide, particularly in the H-4 position, where DIDO1 prefers bulky aromatics, and UL48 requires valine (V) or isoleucine (I) as well as a restricted set of amino-acid residues at the H-3 position. Our observed general preference for hydrophobic residues in the H-4 position is also consistent with the fact that many HCF1-binding peptides have a buried hydrophobic residue in that position (**Fig. 1H, S1A, 2B-C**). Overall, our data strongly suggest that additional molecular determinants of HCF1 binding are encoded beyond the currently recognized consensus “anchor” residues, particularly in the intermediate ‘x’ residue as well as in the flanking N- and C-terminal residues of HCF1-binding peptides.

Next, to further explore this specificity diversity at different positions of HCF1-binding peptides, we further scrutinized our computational HCF1-peptide structural models. These findings point to a dual mechanism of molecular recognition, with certain positions being positively selected for residues that form productive contacts, whereas others are primarily shaped by negative selection against residues that introduce steric hindrance, thereby restricting the permissible sequence space. In our DIDO1 peptide model, we see that residues at, not only its “anchor” aspartate (D), histidine (H) and tyrosine (Y), but also its N-terminal H-4 position, form specific key contacts with HCF1, which suggests that selectivity in these positions is driven by these productive contacts (**Fig. 2D**). Conversely, the selectivity that DIDO1 exhibits at its intermediate ‘x’ and C-terminal Y+1 residues appears most likely to be governed by the need to avoid steric hindrance that any bulkier or physicochemically distinct amino-acid residues would cause (**Fig. 2D**).

To orthogonally validate whether these residues outside of the previously recognized “anchor” positions are *bona fide* molecular determinants of HCF1 binding, we next performed competitive fluorescence polarization (FP) experiments so that we could more directly quantify HCF1-peptide binding affinity differences (**Fig. S2F**). In addition to wild-type DIDO1, DIDO1^WT^, and its non-binding control, DIDO1**^DHxA^**, we tested single point variants at three of these candidate molecular determinants of binding, namely DIDO1**^W^**^-4**S**^, DIDO1**^N^**^x**E**^, and DIDO1**^N^**^+1**K**^ (**Fig. 2C**). In our FP experiments we saw that mutations in these previously unrecognized candidate molecular determinants of binding significantly reduced binding affinity to the Kelch domain of HCF1 (**Fig. 2E, Fig S2G**), with the measured Kd values broadly correlating with our DMS enrichment scores (**Fig. S2H**). These experiments, thus, further support these residues as *bona fide* molecular determinants of binding and propose that HCF1 binding specificity is encoded in an extended sequence of amino-acid residues beyond the previously described “anchor” positions (**Fig. S2I**).

### An extended set of molecular determinants of binding more accurately captures the HCF1 binding landscape

To evaluate the broader relevance of our newly identified molecular determinants of binding in the context of natural HCF1-binding peptides, we scored our screened D/EHxY proteome-wide library using position-specific scoring matrices (PSSM) obtained from our DMS and proteome-wide library experiments. This analysis revealed that functional HCF1-binding peptides have, on average, higher predicted binding scores than the non-binding D/EHxY-containing peptides (**Fig. 3A**), strongly suggesting that incorporating these additional molecular determinants of binding positively contributes to our ability to predict and model HCF1 binding. Moreover, our analysis also predicted E2F1 peptide (together with some other published HCF1-binding peptides) as encoding sub-optimal “non-anchor” residues, which we hypothesized could explain our inability to detect E2F1 binding (**Fig. 1E-F, 3A**).

**Figure 3.**
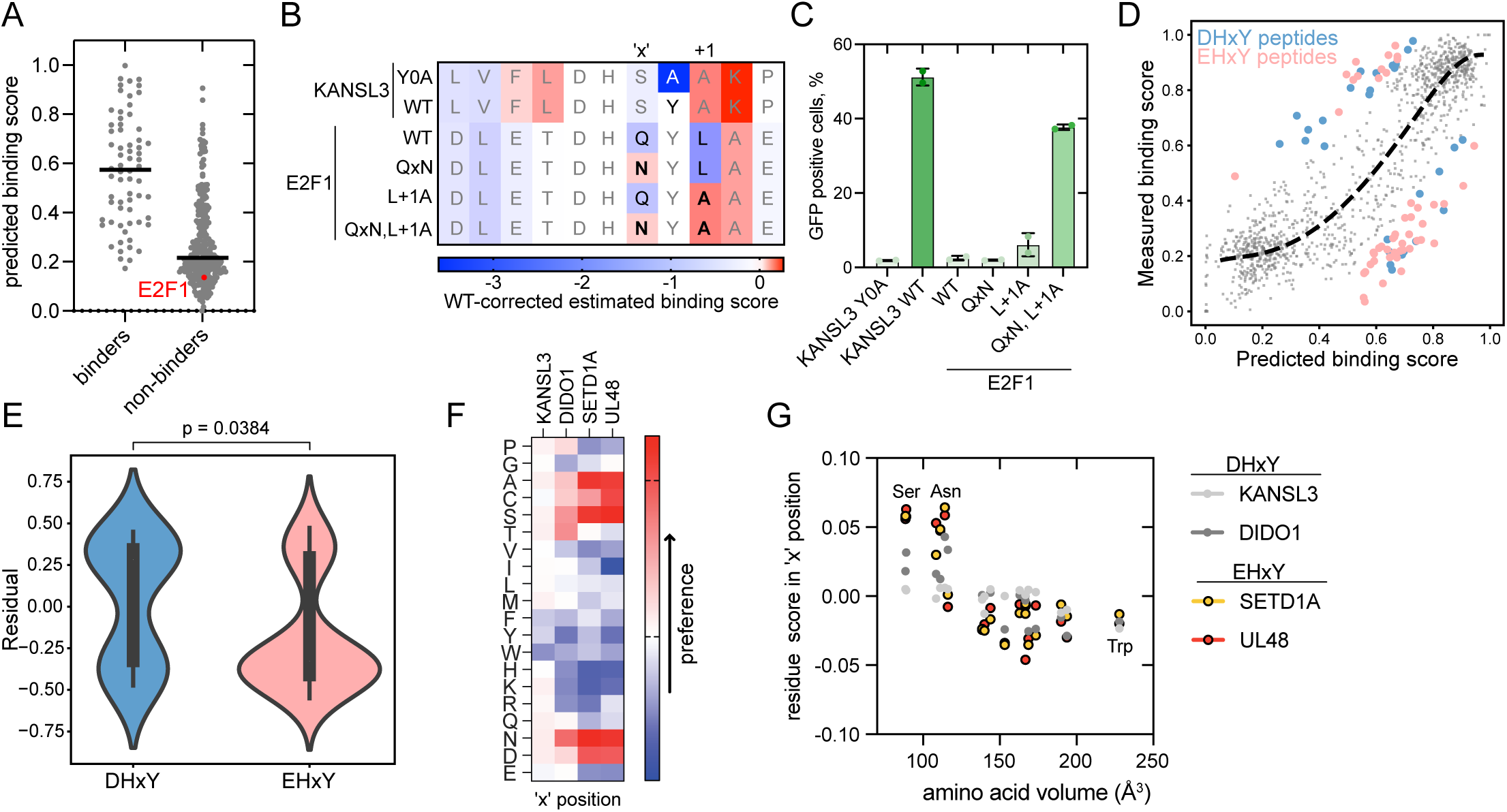
Context specificity and the importance of non-core positions in HBM peptides. (**A**) Scoring of all D/EHxY peptides in the intrinsically disordered regions of the human proteome and validated HCF1 binding motifs with a position-specific scoring matrix obtained from the binding scores of natural (Fig. 1) and DMS peptides (Fig. 2). (**B**) Heatmap showing the residue in each position of E2F1 and KANSL3 peptides with its effect on WT-corrected binding score based on average DMS of two DHxY peptides, DIDO1 and KANSL3. (**C**) Staining of *E. coli* cells displaying the indicated peptides with HCF1(Kelch)-EGFP in flow cytometry. Data is the average of two biological replicates, error bars show standard deviation. (**D**) Scatter plot comparing the enrichment scores predicted by the first-order additive model from PSSM scores to the experimentally measured enrichment score, for each variant. The dotted curve is a monotonic I_spline regression fit from^53^ to match the predicted and observed values, while accounting for global non-linearities. Variants with residuals that deviate from a residual threshold of 2 standard deviations from the best-fit line are considered epistatic, with points above the fit labelled as positively epistatic, and conversely, negatively epistatic below the spline-fit. (**E**) Comparison of the epistasis effect between DHxY and EHxY-type peptides identified from the scatter plot. Statistical significance between the two groups was assessed using a two-sided Mann–Whitney U test, and the corresponding p-value is shown above the plot. (**F**) Heatmap showing the preference of indicated residues in the ‘x’ position in the DMS experiments in Fig. 2C. (**G**) The correlation between residue size and preference score obtained from the DMS experiments in the ‘x’ position of four HBM peptides.

More specifically, based on the DMS data from our four HCF1-binding peptides (**Fig. 2C**), we predicted the E2F1 peptide to encode sub-optimal residues in both its intermediate ‘x’ and its flanking C-terminal Y+1 residues (**Fig. 3B**). To test if the molecular determinants of binding that we obtained from our DMS screens in other HCF1-binding peptides would apply to the E2F1 peptide sequence background, we substituted these intermediate ‘x’ and flanking C-terminal Y+1 residues for residues that we predicted would enhance HCF1 binding. While, as far as our assay was able to detect, the E2F1**^Q^**^x**N**^ mutation did not by itself turn E2F1 into a binder, the E2F1**^L^**^+1**A**^ mutation did enhance HCF1 binding (**Fig. 3B, S3A**), and introducing both of these mutations together in the E2F1**^Q^**^x**N,L**+1**A**^ variant enhanced HCF1 binding to a level comparable to KANSL3, one of the tightest natural binders of HCF1 (**Fig. 3B**). Thus, these experiments show that the rules learned from the DMS of four peptides apply to other HCF1 binding peptides and that our newly identified molecular determinants of binding are able to enhance HCF1 binding.

To further examine the impact of peptide sequence background on positional preferences, we investigated the correlation between measured and predicted binding scores among both natural (**Fig. 1**) and DMS peptides (**Fig. 2**). Through this analysis, we observed that a small subset of peptides overperformed their predicted binding scores, while a separate subset of peptides underperformed them (**Fig. 3C**). Our finding that 75% (36 out of 48) of all underperforming peptides encoded glutamate (E) at the H-1 position (**Fig. 3D-E, S3B**), suggested to us that encoding this amino-acid residue at this position may be negatively coupled or, in other words, decrease the possible and preferred residues, in other positions. Also consistent with this hypothesis is the fact that our DMS results from the two DHxY libraries and two EHxY libraries showed stronger selectivities in the intermediate ‘x’ position of the EHxY peptides, where we observed a more significantly pronounced preference for small residues (**Fig. 3F-G**). Altogether, our experiments suggest that while most HCF1-binding peptides have had to evolve under similar positional residue constraints, some context-specific effects, such as encoding a glutamate residue (E) at the H-1 position, have likely driven coupled constraints at other positions.

### The natural HCF1-binding landscape extends to proteins with non-canonical, register-shifted two-residue sequences spanning their anchor histidine and tyrosine residues

Given the diversity we uncovered around its previously established binding sequences, we next asked if the Kelch domain of HCF1 could also recognize and bind peptides encoding sequences beyond its consensus D/E/N/QHxY sequences. To this end, we investigated all HCF1-binding proteins that we had previously identified as being outcompeted by a DHxY-containing peptide (**Fig. 4A**)^11^. While out of the 101 outcompeted proteins, 32 contained a D/E/N/QHxY sequence and 52 were known interactors of these D/E/N/QHxY-containing proteins (thus nominating them as likely indirect HCF1 interactors), 18 did not fulfill either condition (**Fig. 4A, S4A**). Computational AlphaFold3 modeling of these 18 non-canonical candidate HCF1-binding proteins led us to narrow down to the only candidate that AlphaFold3 was able to predict with high confidence (ipTM > 0.8)^26^, SMCHD1 (**Fig. S4B**). When comparing this SMCHD1 model with DIDO1, we observed that the SMCHD1^214DHSGY218^ peptide is bound to HCF1 in a conformation where the “anchor” aspartate (D), histidine (H) and tyrosine (Y) residues effectively align with the analogous residues in the DHxY DIDO1 peptide. However, we simultaneously noticed a critical difference in the extended, register-shifted two-residue sequence spanning its anchor histidine (H) and tyrosine (Y) residues in SMCHD1 (**Fig. 4B**). Thus, while several molecular determinants of binding for HCF1 appear to be conserved among many of its binders, at the same time, there appears to be a more diverse and non-canonical landscape of HCF1 binders than previously recognized.

**Figure 4.**
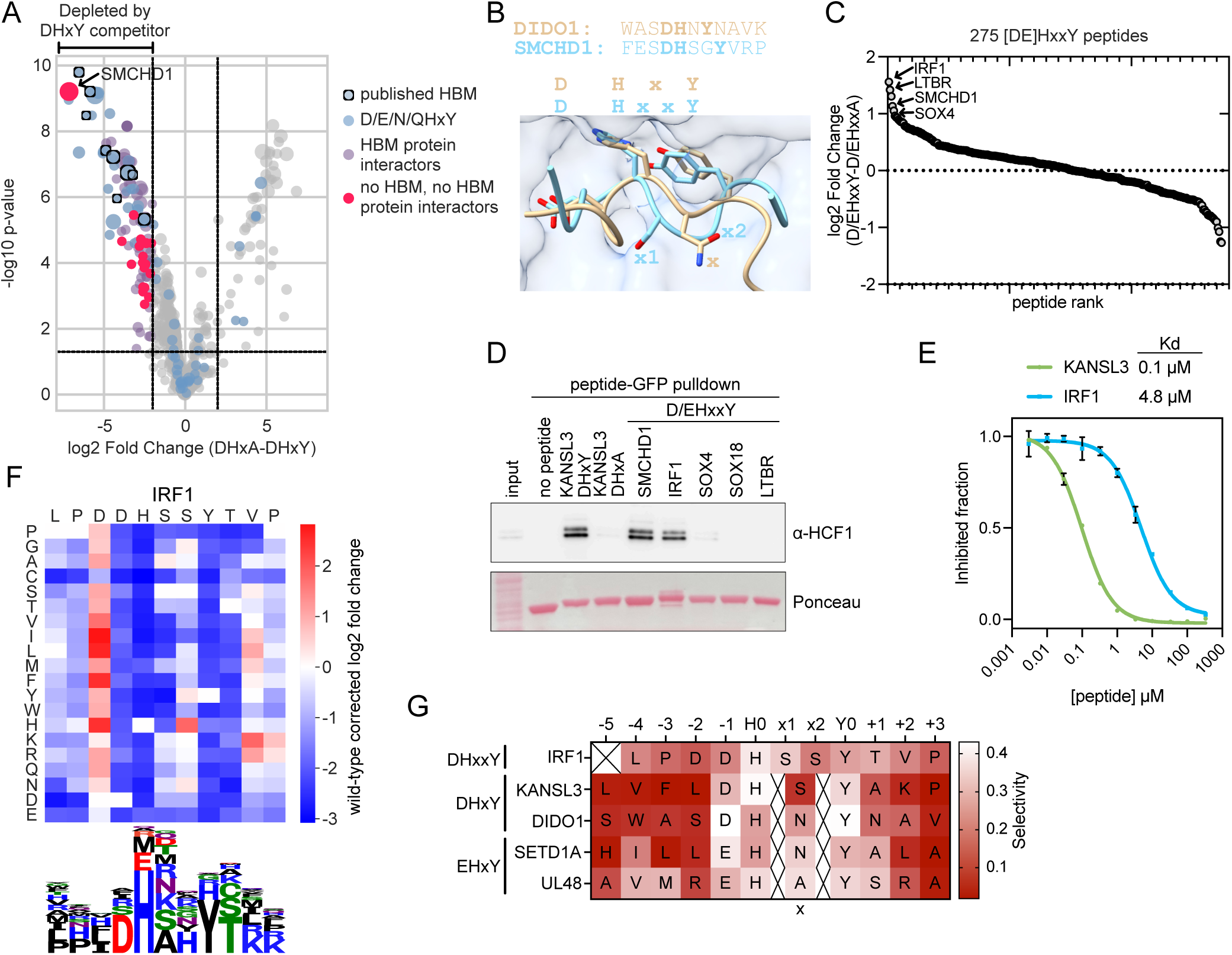
HCF1 interacts with unconventional DHxxY-type motifs. (**A**) Plot showing the proteins outcompeted in HCF1 binding by KANSL3 DHxY competitor peptide. Data is from^11^. Mapping of HCF1 binding motifs and the protein interaction network is in Fig. S4A. (**B**) Superimposed DIDO1 DHxY and SMCHD1 DHxxY peptides from peptide-HCF1 complex AlphaFold3 models. (**C**) HCF1(Kelch)-GFP binding of 275 D/EHxxY peptides from the human proteome measured in bacterial surface display. (**D**) Peptide-GFP pulldown from HCT116 cell lysate using recombinant peptides to test binding to HCF1. A representative example of two biological replicates is shown. (**E**) Dose-response curves measuring KANSL3 and IRF1 peptides as competitors to displace a FITC-labeled HCF1 binding peptide from HCF1, measured by fluorescence polarization (n = 2 biological replicates). (**F**) Wild-type-corrected HCF1 binding scores measured in bacterial peptide display with DMS libraries of IRF1 peptide. Below, the sequence logo showing positional preferences for HCF1 binding to IRF1. (**G**) The Gini index (selectivity) showing inequality between residues in a position, calculated from DMS data, and the number of van der Waals contacts of the residue with HCF1 in AlphaFold3 model.

To further evaluate the degree to which such non-canonical binding may exist for other natural HCF1 binders, we next asked if additional DHxxY-containing peptides exist in other natural proteins and whether they could also bind HCF1. To this end, we used SLiMSearch^27^ to identify candidate HCF1 binders containing this sequence from the intrinsically disordered regions of the proteome, which revealed 290 candidates. To determine if these peptides bind HCF1, we carried out a bacterial surface display screen including these 290 D/EHxxY-containing peptides alongside their counterpart DHxxA non-binding control peptides. While compared to the proteome-wide screen of conventional D/EHxY peptides this D/EHxxY peptide library led to lower enrichment scores likely reflecting weaker binding overall (comparing **Fig. 1G** and **S4D**), we identified peptides from several transcriptional regulators as enriched in this D/EHxxY-focused screen (**Fig. 4C, Table S1**). To further validate these candidate HCF1-binding proteins from our proteome-wide screen as *bona fide* HCF1 binders, we performed peptide pulldown experiments, which provided further evidence that the DHxxY peptides from SMCHD1 and IRF1 bind HCF1 (**Fig. 4D**). Through follow-up fluorescence polarization assays, we estimated the dissociation constant of the IRF1 peptide to be around 4.81 μM while for the DHxY-containing KANSL3 and DIDO1 we estimated them to be around 0.1 μM and 1.15 μM, respectively (**Fig. 2E and 4E**). While these dissociation constant differences together with the fact that IRF1 is among the most enriched within our DHxxY-focused proteome-wide library suggest that this non-canonical binding mode may yield weaker affinity than the canonical D/EHxY sequence, the IRF1 DHxxY peptide binds HCF1 with an affinity comparable to several of D/EHxY peptides (**Fig. 2E**).

To further elucidate the molecular determinants of binding for these non-canonical HCF1 binders, we performed DMS screening of the IRF1 peptide and evaluated HCF1 binding using our bacterial peptide display assay (**Fig. S4D**). Analogous to our DMS screen results in the background of D/EHxY peptides, we observed the strongest selectivity in the “anchor” aspartate (D), histidine (H) and tyrosine (Y) residues, where no other residues were tolerated (**Fig. 2C** and **4F**). These results further support the view that these “anchor” residues are configured and bind similarly in both canonical and non-canonical HCF1 binders – a view that is also supported by our AlphaFold3 models (**Fig 4B**). However, while for our canonical HCF1 binders glutamate (E) is tolerated at the H-1 position, for the non-canonical IRF1 there is a clear preference for aspartic acid at this position. In line with our observations in our D/EHxY DMS screens, we here find that additional molecular determinants of binding are encoded beyond the “anchor” aspartate (D), histidine (H), and tyrosine (Y) residues, in the intermediate “x1” and “x2” residues as well as in the N- and C-terminal flanks of this non-canonical peptide. Whereas the first intermediate “x1” position has a moderate preference for alanine and disfavors the larger hydrophobic residues, isoleucine (I) and leucine (L), and aromatic residues, the second intermediate “x2” position shows a preference for histidine (H), while being more tolerant to other residues than the preceding intermediate “x1” position. Moreover, we also observed selectivity at the N- and C-terminal flanks of the peptide, particularly in the Y+1 and Y+2 positions, where IRF1 strongly disfavors proline (**Fig. 4F**), similarly as we had observed for D/EHxY peptides (**Fig. 2C**). To more directly compare the specificity of D/EHxY and DHxxY peptides, we calculated selectivity scores for each DMS position in the sequence backgrounds of KANSL3, DIDO1, SETD1A, UL48, and IRF1. Compared to KANSL3, IRF1 shows stronger selectivity in its intermediate “x1” and “x2” position(s) and in its N- and C-terminal flanks which, in turn, makes its specificity more comparable to that of UL48 (**Fig. 4G**). Thus, our DMS screen results in the IRF1 peptide sequence background further support this non-canonical DHxxY sequence as a *bona fide* HCF1 binder that, while sharing some features with the previously recognized D/EHxY binding sequence, also encodes additional molecular determinants of binding elsewhere.

### The non-canonical DHxxY sequence promotes the transcriptional activity of IRF1, which can be enhanced by potentiating its binding affinity for HCF1

IRF1 is a transcription factor that becomes activated in response to pathogens, promotes G1 cell cycle arrest and inhibits cell proliferation^28,29^. To test if the ^160^DHSSY^164^ in IRF1 is required for its molecular activities, we induced the expression of either wild-type IRF1, IRF1^WT^, the tyrosine-to-alanine HCF1-non-binding negative control mutant, IRF1^NB^, or an IRF1 mutant where we replaced its DHxxY sequence with a high-affinity HCF1-binding sequence grafted from KANSL3 (LVFLDHSYAKP, **Fig 4E, 5A**), IRF1^HB^, in MDA-MB-231 breast cancer cells. Compared to the non-binding IRF1^NB^ control, we observed that IRF1^WT^ had an anti-proliferative phenotype, which was further increased in the high-affinity HCF1 binder IRF1^HB^ variant (**Fig. 5A, S5A**). Similarly, while the expression of the non-binding negative control IRF1^NB^ did not perturb the cell cycle, expression of IRF1^WT^ and grafted IRF1^HB^ resulted in a moderate and more pronounced accumulation of cells in G1, respectively (**Fig. 5B**). The substantially decreased phenotypic effects that we observed in our IRF1^NB^ mutant suggest that the DHxxY sequence is required for its cell cycle and anti-proliferative molecular functions. The fact that IRF1 activity can be enhanced by grafting a stronger HCF1-binding sequence further supports that the effects we observe are largely driven by HCF1 binding and that the IRF1 ability to bind HCF1 may be an important driver of its downstream transcriptional activity.

**Figure 5.**
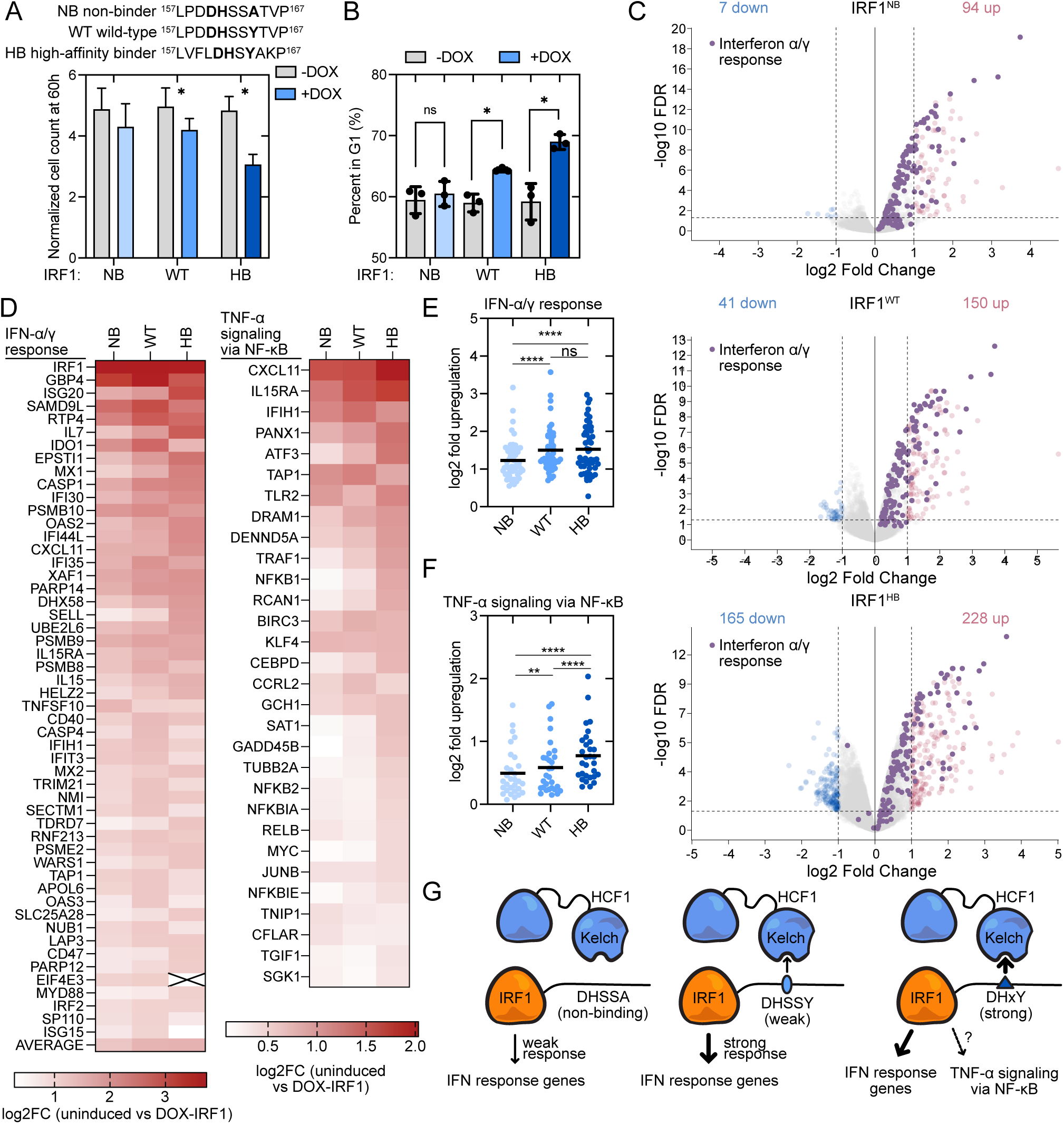
DHxxY motif is necessary for full transcriptional activity of IRF1. (**A**) The effect of overexpression of different IRF1 variants on MDA-MB-231 cell proliferation. In IRF1-NB, the DHxxY motif is inactivated with Y164A mutation. In IRF1-HB, the DHxxY motif is replaced with a stronger DHxY motif from KANSL3. Data is average from three biological replicates, error bars show standard deviation. (**B**) The effect of different IRF1 variants on cell cycle measured by DNA staining using propidium iodide. Data is average from three biological replicates, error bars show standard deviation. (**C**) RNAseq experiment showing the effect of doxycycline-driven induction of wild-type and mutant IRF1 on gene expression. (**D**) The impact of different IRF1 variant overexpression on interferon response genes and genes involved in TNFα signalling via NF-kB. (**E**) Upregulation of interferon response genes by different IRF1 variants measured in RNAseq. (**F**) Upregulation of TNFα signalling via NF-kB genes by different IRF1 variants measured in RNAseq. (**G**) Model of IRF1 co-regulation by HCF1.

To obtain a more detailed understanding of the role of HCF1 binding in driving the transcriptional function of IRF1, we performed RNAseq on cells overexpressing our different IRF1 variants. These experiments showed that inducing IRF1 expression drives upregulation of genes involved in interferon α/ɣ signaling (**Fig. 5C, S5B, Table S3**) which are known targets of IRF1^30,31^. While we also observed an upregulation of the interferon signaling proteins in cells expressing the HCF1-non-binding negative control IRF1^NB^ mutant, this transcriptional response was significantly blunted compared to wild-type IRF1 (**Fig. 5D-E**), suggesting that while the HCF1 interaction may not be essential for part of the activity of IRF1, HCF1 binding does contribute to IRF1 transcriptional activity. While IRF1^WT^ and IRF1^HB^ activated interferon signalling genes similarly (**Fig. 5E**), the IRF1^HB^ variant drove a broader transcriptional response, upregulating 228 genes (**Fig 5C**), and simultaneously promoting an upregulation of genes involved in TNFα signaling through NFkB (**Fig. 5F, S5C-D**). Supporting the physiological relevance of these findings, IRF1 is known to promote NFkB gene transcription and form a complex with NFkB to activate its downstream targets in inflammatory signalling^32,33^. Together, these data show that the non-canonical HCF1-binding sequence in IRF1 is required for its full activity and suggest that HCF1 binding affinity is an important contributor to the signaling output of HCF1 partners, such as IRF1 (**Fig. 5G**).

### Canonical and non-canonical HCF1-binding proteins become O-GlcNAcylated

Given that HCF1 has been found to mediate O-GlcNAcylation of its Kelch domain binders NRF1 and PGC1ɑ, and many proteins involved in transcription have been found to be O-GlcNAcylated^7,8,32–34^, we next set out to examine if HCF1 interaction could be a widespread mechanism promoting O-GlcNAcylation. With the aim to quantify whether a significant fraction of HCF1 binders are O-GlcNAcylated, we overexpressed the HCF1-interacting KANSL3 peptide fused to GFP in HCT116 cells to outcompete endogenous HCF1 interactions, and performed a coIP using an antibody against O-GlcNAc modification, which we analyzed by mass-spectrometry. These experiments revealed that 39 proteins are less O-GlcNAcylated upon expression of DHxY competitor peptide (**Fig. 6A, Table S4**) strongly suggesting that they are substrates that become O-GlcNAcylated upon HCF1 binding. The fact that the majority of these (17 out of 39) contain HCF1 binding peptides and that specific O-GlcNAcylation sites have been previously identified in 23 of them^35^, further supports that the majority of them are directly O-GlcNAcylated (**Fig. 6A**). Among these HCF1-binding candidate O-GlcNAcylation targets, we observed enrichment in histone modifiers and transcription regulators (**Fig. 6B**).

**Figure 6.**
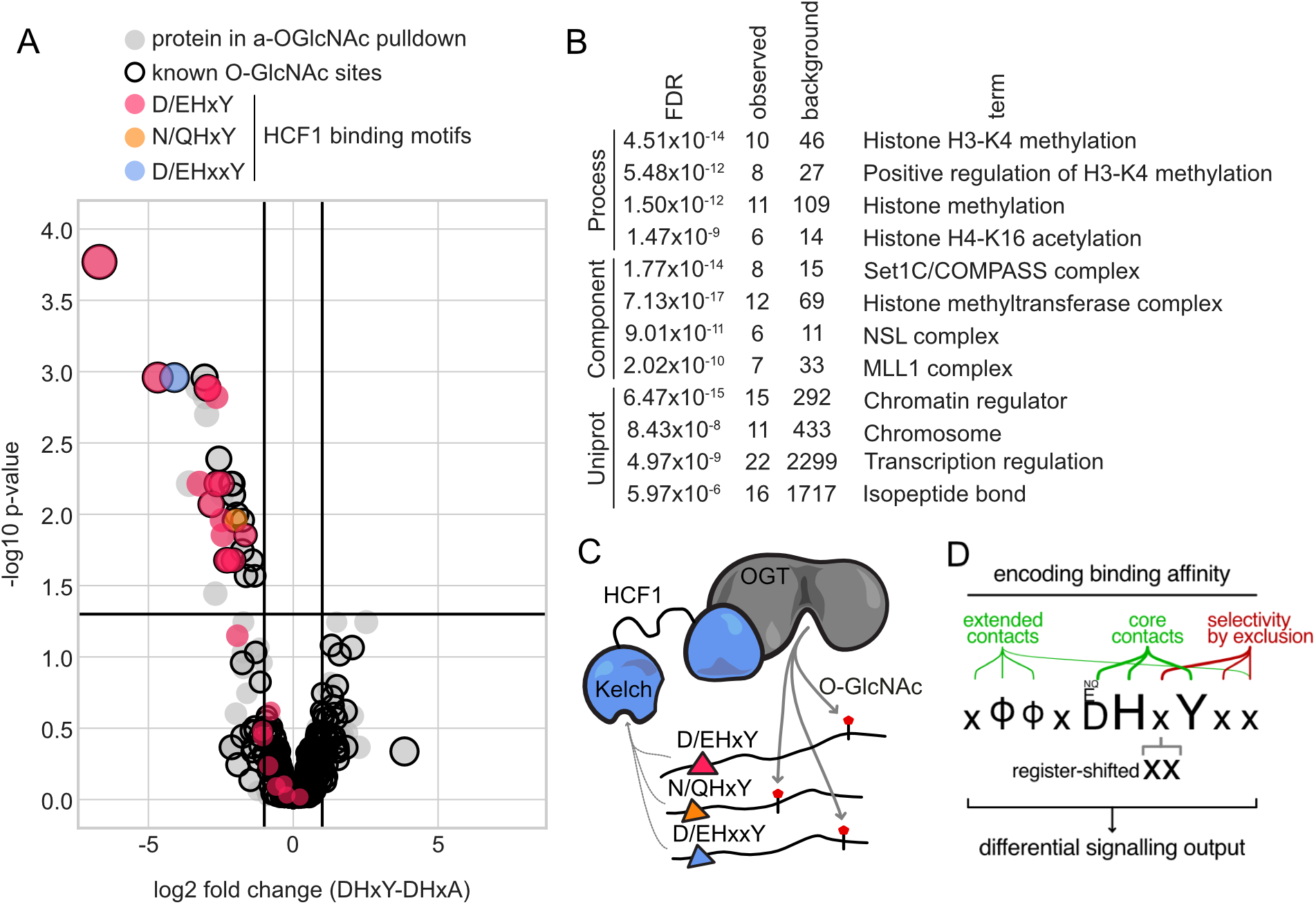
HBM interaction mediates O-GlcNAcylation of nuclear proteins. (**A**) Vulcano plot showing decrease in O-GlcNAc-modification of 39 proteins upon expression of KANSL3 DHxY competitor peptide in HCT116 cells. (**B**) Enrichment of GO terms (Process, Component, and UniProt keywords) among the 39 proteins that are less O-GlcNAc modified in the presence of DHxY competitor peptide in HCT116 cells. (**C**) Scheme illustrating the role of Kelch domain of HCF1 in recruiting glycosylation substrates for OGT. (**D**) Summary model of HCF1 Kelch domain docking interactions. Φ denotes hydrophobic residues.

Together, our results suggest that HCF1, by interacting with a large number of proteins containing both canonical and non-canonical HCF1-binding sequences, promotes the O-GlcNAcylation of transcriptional proteins by effectively acting as a substrate recruitment scaffold to the O-GlcNAcylation enzyme, OGT (**Fig. 6C**). Given that the modification of many proteins is not affected by the HCF1-binding competitor peptide (**Fig. S6A**), we also conclude that OGT has other substrate recruitment mechanisms. Even though OGT is localised both in the nucleus and cytoplasm^36^, the fact that 36 out of the 39 proteins with decreased detection in our O-GlcNAc coIP experiments are nuclear (**Table S5**), also suggests that HCF1 contributes and promotes O-GlcNAcylation of nuclear targets.

## Discussion

The evolution of protein interacting domains is often constrained by the need to interact with a large number of binding partners using a single molecular pocket. In light of this, it is through the systematic characterization of molecular determinants of binding that we become more broadly knowledgeable and capable of predicting and testing novel candidate binders, evaluating relative binding strengths, and ultimately shed light onto the molecular functions that proteins play. On this note, our proteome-wide experiments testing peptides with predicted canonical HCF1-binding sequences allowed us to identify 41 previously uncharacterised peptides as HCF1 binders (**Table S1**), including eight transcription factors (MXD3, CREB3L1, FOXN3, ZBTB38, ZNF704, LMX1A, TBX21, MEIOSIN) and five chromatin regulators (KMT2B, PHF2, TASOR2, BRWD3, ZNF518A), thereby further expanding the known and widespread role of HCF1 in transcription. We also find HCF1-binding sequences in proteins involved in SUMOylation (USPL1, SENP5)^37^, a PP2A regulatory subunit (PPP2R2B) and proteins associated with E3 ubiquitin ligase NEDD4-1 (N4BP2, BEAN1). In addition to the known canonical D/E/N/QHxY HCF1-binding sequence, we find that HCF1 binds non-canonical DHxxY peptide sequences in an epigenetic regulator, SMCHD1, and a transcription factor, IRF1 (**Fig. 4**). Consistent with our findings for HCF1, and further supporting their physiological relevance, the paralog of HCF1, HCF2, has been found to bind IRF1 through its Kelch domain and potentiate the activity of IRF1 in response to viral infection^38^. The fact that from the 863 canonical (D/E/N/QHxY) and non-canonical (D/EHxxY) candidate HCF1-binding sequences, we only found 69 to be likely *bona fide* HCF1 binders (**Table S1**) highlights the importance of characterizing molecular interactions in greater depth and beyond previously recognized binding sequences and molecular determinants of binding.

As a case in point, our experiments uncovered key contributions from non-core residues in both driving whether a peptide will bind HCF1 at all and in fine-tuning its binding affinity. Similar principles have been recognized in other peptide-domain interactions^25,39–42^. By mutating and grafting into its HCF1-binding sequence, we found that differential binding to HCF1 critically contributes to the transcriptional output of IRF1, highlighting the role of binding affinity in HCF1-mediated transcriptional co-regulation (**Fig. 5**). We find two major mechanisms driving specificity in “non-anchor” residues (**Fig. 6D**). First, we find a specificity driven by residues, such as DIDO1^W505^ in the H-4 peptide position, that appear to make additional contacts with the HCF1 pocket. In addition, we find a second type of specificity, such as the selectivity in the intermediate ‘x’ and C-terminal Y+1 positions, which appears to be driven by avoidance of steric hindrance, indicated by preferences for small residues (instead of for similar residues that could form similar contacts). The importance of “non-anchor” positions varies between HCF1-binding proteins. While we observed an overrepresentation of DHxY (51 out of 66) over EHxY peptides (15 out of 66) among HCF1 binders in our proteome-wide library, this H-1 aspartate-to-glutamate bias cannot be fully explained by overall intolerance or residue preference, as aspartate (D) is only preferred over glutamate (E) in one of the four peptide context backgrounds we tested (**Fig. 2C**). We also identified a relatively smaller fraction of N/QHxY and DHxxY non-canonical peptides as HCF1 binders compared to canonical D/EHxY peptides, even though the functional DHxxY peptides that we identified bind HCF1 with comparable affinity as the canonical peptides (**Fig. 2E, 4D**). These observations, together with our DMS data, put forward a model where selectivity in “non-anchor” positions may function as a key driver in determining what may be a predominant binding mode for a given molecular pocket.

Despite binding to over 50 transcription factors and regulators, the molecular mechanisms around how HCF1 may contribute to transcription have remained elusive. Together, and consistent with previous studies^7,8^, we propose that HCF1 functions as a mediator of O-GlcNAcylation by bringing proteins containing canonical and non-canonical HCF1-binding sequences to the proximity of OGT. O-GlcNAcylation has been shown to modulate the function of transcription factors and other proteins through multiple mechanisms including disrupting protein-protein interactions^43^, regulating protein stability^44^, promoting nucleo-cytoplasmic shuttling^45^, and as well as more directly contributing to transcriptional activity^46^. Given that similar to other post-translational modifications, O-GlcNAcylation can act both positively or negatively on its downstream regulated proteins depending on the specific protein target in question, its direct link to this post-translational modification could also help explain some of the context-specific effects of HCF1^18,47^. HCF1 and O-GlcNAcylation have been found to promote cancer progression, with various cancers showing increased O-GlcNAcylation levels^14,48,49^. In a recent study, we found that despite HCF1 being a common essential gene, different cancer cell lines presented variable sensitivity to the Kelch pocket inhibition^11^. Consistent with our results and those of others, inhibiting the Kelch domain of HCF1 has been proposed as an approach to suppress MYC-driven cancers^50^. Altogether, given the link between O-GlcNAcylation and cancer and the large number of oncogenes that bind to HCF1, further studies into the therapeutic potential of HCF1 are warranted.

## Methods

### Cell culture

HCT116 cells were obtained from ATCC. MDA-MB-231 cells were a kind gift from J. Carroll lab at Cancer Research UK Cambridge Institute. HCT116 was cultured in McCoy 5A medium (36600021, Gibco), MDA-MB-231 in ATCC-modified RPMI 1640 medium (11504566, Gibco), and HEK293T in DMEM with pyruvate (41966029, Gibco). All media were supplemented with 10% fetal bovine serum (A5209402, Gibco), 100 U/ml penicillin, and 0.1 mg/ml streptomycin (15140122, Gibco). Doxycycline hyclate (D9891-1G, Sigma-Aldrich) was dissolved at 1 mg/ml in DMSO. All cell lines were cultured in a 37 °C incubator with 5% CO_2_. All cell lines were tested negative for Mycoplasma by qPCR at the Cancer Research UK Cambridge Institute Research Instrumentation and Cell Services core facility.

The competitor peptide and EGFP coding sequences as well as IRF1 and HCF1(Kelch) constructs were cloned into doxycycline-inducible lentiviral vector pCW57.1 (Addgene #41393) using NEB Stable Competent *E. coli* cells, followed by plasmid DNA extraction with QIAfilter plasmid kits (Qiagen). The virus was produced as follows: 7 × 10^5^ HEK293T/17 cells (ATCC: CRL-11268) were seeded to 6-well plate and in 24 h were transfected using lipofectamine 3000 (Thermo Fisher Scientific) with psPax2 (Addgene #12260) and pMD2.G (Addgene #12259) as packaging plasmids. 6 h after the transfection, the medium was replaced with 3 ml fresh DMEM. For adherent cell lines, 2-3 × 10^5^ cells were seeded to a 6-well plate 24 h before transduction. The medium containing the virus was collected 2 days after transfection and together with 8 µg/ml polybrene (TR-1003-G, Sigma-Aldrich) was used to transduce the recipient cells. In two days, the transduced cells were split and supplemented with 1 µg/ml puromycin to eliminate untransduced cells. Prior to seeding for experiments, the transduced cells were split at least 3 times and cultured in the presence of 1 µg/ml puromycin (A1113803, Gibco) to allow selection of the transduced cells.

### Strep-tagged HCF1(Kelch) purification

HEK293T cells stabling expressing HCF1(Kelch)-Strep or HCF1(Kelch)-GFP were seeded on 15 cm plates with 1 µg/ml puromycin for selection. When the cells reached 50-60% confluency, protein expression was induced with 1 µg/ml doxycycline. Following 36 hours of induction, the cells were collected by scraping, centrifuged at 300xg for 5 minutes, and washed twice with chilled PBS. Cells were resuspended in lysis buffer containing 20 mM Tris-HCl pH 7.5, 150 mM NaCl, 1 mM EDTA, 1% Triton-X100, 1 mM DTT, cOmplete EDTA-free Protease Inhibitor Cocktail (Roche), a PhosSTOP tablet (Merck), and Biolock Biotin blocking solution (IBA). The lysate was incubated rotating for 20 minutes at 4°C and then pulled through a syringe to aid lysis. Debris was pelleted by centrifugation at 21,000 xg for 15 minutes at 4°C. The supernatant was subsequently loaded on to a column containing 100 µL of Strep-Tactin-XT resin (IBA) which has been equilibrated in Buffer W (IBA). The flowthrough was collected and loaded 3 times on the column to maximize binding, and then the resin was washed with 25 column volumes of Buffer W. The protein was eluted in 8 fractions, the first with 100 µL of Buffer BXT (IBA) and the following fractions with 200 µL Buffer BXT. For the third, fifth, and eighth elutions, the column was capped and the resin was incubated with the buffer for five minutes to maximize yield. The fractions were run on an SDS-PAGE gel to verify their purity before pooling together fractions containing the protein of interest. Concentration of the pooled samples was determined using bovine serum albumin standards (Thermo Fisher Scientific).

### Bacterial peptide display

HBM peptides of interest were cloned to pBAD33-eCPX plasmid resulting in a fusion protein containing an N-terminal OmpX signal peptide, desired HBM, eCPX, and a C-terminal MYC tag. Electrocompetent *E. coli* MC1061 cells were transformed with plasmid and grown overnight at 37°C and 220 rpm in LB with 25 µg/ml chloramphenicol. The cultures were diluted 1:100 and grown to an OD600 of 0.5-0.6 then induced with 0.4% arabinose at 18°C for 16-18 hours. Cells were pelleted by centrifugation at 4,000 x g and 4°C. Pellets were washed with chilled PBS and resuspended to an OD of 1. 100 µl of cells were pelleted and resuspended in 30 µl of staining buffer, 0.2% BSA in PBS with Alexa-647 anti-MYC (Biolegend, 626810) and GFP-tagged HCF1(Kelch) protein. Anti-MYC antibody was used at a 1:30 dilution, and 500 ng of GFP-HCF1 protein was added per sample. Cells were incubated with the staining solution for approximately 45 minutes at 4°C and then washed twice with chilled PBS. Samples were resuspended in 100 µl of PBS and diluted 1:10 before being analyzed by fly cytometry (BD LSR-Fortessa^TM^ Cell Analyser). All data was processed using FlowJo 10.10.0.

For cell sorting, following staining, 1 ml 0.2% BSA in PBS was added to the sample and then immediately sorted on BD Influx System at the University of Cambridge Cambridge Institute for Medical Research Flow Core Facility.

### AlphaFold3 models and analysis

AlphaFold3^26^ was used to predict structures of complexes containing the Kelch domain of HCF1 (positions 1-360) and peptides. HCF1-peptide models where any of the key [DE]HxY residues were oriented towards the solvent not HCF1 were discarded as incorrect. To identify buried residues, the solvent accessible surface areas of each residue of the peptide were calculated in the HCF1-bound and -unbound states using ChimeraX^51^.

### Fluorescence polarisation

Unlabeled peptides and N-terminally FITC-Ahx labeled KANSL3 HBM peptide (VFLDHSYAK) with >95% purity were ordered from GenScript. Fluorescence polarization was measured in a buffer containing 25 mM Hepes-KOH pH 7.4, 150 mM NaCl, 5 mM DTT and 0.05% CHAPS. The assays were carried out in black, low volume, non-binding, 384-well plates (4514, Corning). After mixing, the reactions were incubated at 37 °C in the dark for 30 min before measuring fluorescence polarization using PHERAstar FSX microplate reader. Peptide binding affinities were measured by dosing unlabeled peptides that compete with the FITC-KANSL3 HBM peptide for binding to the Kelch domain of HCF1. In competitive FP experiments, both FITC-KANSL3 HBM peptide and Kelch domain of HCF1 were used at 25 nM concentration. The fluorescence polarization experiments were performed in at least two replicates. The data was analyzed and plotted using Prism 10 (Graphpad).

### Peptide-EGFP-6xHis protein purification

For recombinant protein purification, the peptide-GFP constructs were cloned to pET28a, resulting in C-terminal tagging with 6xHis tag. The protein expression in *E. coli* BL21(DE3) cells was induced at culture optical density OD600=0.6 by addition of 1 mM IPTG, followed by further incubation at 37 °C for 3 h, after which the cells were pelleted and frozen. The cell pellets were resuspended in a buffer containing 50 mM Tris-HCl, pH 7.4, 300 mM NaCl, 1% Triton X-100, 5% glycerol, 10 mM imidazole and cOmplete, EDTA-free Protease Inhibitor Cocktail (Roche), and 1 mg/ml lysozyme (Sigma-Aldrich). The lysate was incubated on ice for 15 min, sonicated, and cleared by centrifugation at 4 °C 20 000 g for 15 min. The 6xHis-tagged proteins were bound to HisPur™ Ni-NTA Resin (Thermo Fisher Scientific) in gravity flow columns and washed 4 times with the lysis buffer supplemented with 25 mM imidazole. The proteins were eluted with 50 mM Tris-HCl, pH 7.4, 500 mM NaCl, 0.5% Triton X-100, 5% glycerol, and 200 mM imidazole. The eluates were analyzed by SDS-PAGE to verify the purity and to evaluate protein concentration using bovine serum albumin standards (Thermo Fisher Scientific).

### Peptide-GFP pulldown experiments and O-GlcNAc coIP

To test the interaction between recombinant HBM peptides and HCF1, the peptide-GFP proteins were loaded onto ChromoTek GFP-Trap® Magnetic Agarose (Proteintech), followed by incubation with human cell lysate containing endogenous HCF1.

HCT116 cells were grown in 15 cm plates to 80% confluency, when they were collected by scraping and frozen. The cells were lysed in buffer containing 10 mM Tris-HCl, pH 7.4, 140 mM NaCl, 10% glycerol, 0.5% NP-40, 0.25% Triton X-100, 1 mM EDTA, 1 mM DTT and cOmpleteTM protein inhibitors cocktail (Merck). The lysates were pulled through a 26G needle and cleared by centrifugation at 20 000 g 4 °C 10 min. The total protein concentration in the lysate was measured using Pierce™ Bradford Plus Protein Assay Reagent (Thermo Fisher Scientific). The lysates were diluted to 5 mg/ml total protein. For peptide-GFP pulldown, 3 µg of peptide-EGFP protein was loaded onto 5 µl ChromoTek GFP-Trap® Magnetic Agarose beads (Proteintech) that had been equilibrated with the lysis buffer. Unbound peptide-GFP was washed off the GFP-Trap agarose beads, and 1-2 mg of cell lysate was mixed with the peptide-GFP loaded beads. The mixture was incubated on a rotator at 4 °C for 1 hour. Then, the GFP-trap magnetic beads were collected using a magnet, the lysate was removed and the beads were washed three times with 1 ml lysis buffer. The immunoprecipitated proteins were eluted using Laemmli SDS-PAGE sample buffer and were subjected to SDS-PAGE followed by immunoblotting.

For the a-O-GlcNAc coIP, HCT116 cells were treated with 1 µg/ml doxycycline to induce expression of wild-type KANSL3 HBM or its negative control peptide-GFP fusions for 24h before collecting. The cells were lysed as described above, followed by lysate preclearing by incubating on a rotator with 25 µl protein A Dynabeads (10002D, Invitrogen) for 1 hour at 4 °C. Then, the supernatant was mixed with 1 µg O-GlcNAc MultiMab® Rabbit mAb mix (#82332, Cell Signaling Technologies) bound to 25 µl protein A Dynabeads (10002D, Invitrogen) to precipitate O-GlcNAc-modified proteins from HCT116 cell lysate. The lysate was incubated with the antibody-bound beads overnight at 4 °C. The beads were washed three times with the lysis buffer, followed by two washes with 100 mM ammonium bicarbonate, moving the beads to a new tube with each wash. Further sample processing, including trypsinisation to peptides and peptide purification, and mass-spectrometry with Orbitrap Fusion Lumos Tribrid Mass Spectrometer (Thermo Fisher Scientific) was performed at the Cancer Research UK Cambridge Institute Proteomics core facility. Functional enrichment analysis of outcompeted proteins was performed with STRING^52^.

### Immunoblotting

For immunoblotting, the cell pellets were resuspended in a buffer containing 10 mM Tris-HCl pH 7.4, 150 mM NaCl, 1% Triton X100, phosSTOP phosphatase inhibitor cocktail (Roche) and cOmplete EDTA-free protease inhibitor cocktail (Merck) and pulled through a 26G needle. The lysate was cleared by centrifugation and the total protein concentration was measured using Pierce™ Bradford Plus Protein Assay Reagent. For immunoblotting, 20-40 µg of cell lysate was resolved using SDS-PAGE and the proteins were transferred to nitrocellulose membrane using iBlot 3 (Thermo Fisher Scientific). After transfer, the membrane was blocked using 5% fat-free milk solution in TBS-T, followed by overnight incubation at 4 °C with the primary antibody solutions. Hcfc1 Antibody (Amino-terminal Antigen) (#69690, Cell Signaling Technology) was used at 1:1000. Then, the membrane was washed 5 times with TBS-T, incubated with secondary antibody solutions for 1 hour, and washed again 5 times with TBS-T. HRP-conjugated anti-rabbit IgG (#7074, Cell Signaling Technologies) was used at 1:10000. The antibodies were detected using SuperSignal™ West Pico PLUS Pico Chemiluminescent substrate (Thermo Fisher Scientific) and Amersham ImageQuant.

### RNAseq

MDA-MB-231 transduced with doxycycline-inducible IRF1 were cultured to 40% confluency in 6-well plates and treated with 0.05 µg/ml doxycycline for 24h to induce IRF1 overexpression. The cells were harvested by trypsinization, pelleted and resuspended in Zymo DNA/RNA Shield buffer (Zymo Research). Cells were diluted to 200K cells/ml and 100K cells were submitted to bulk RNA sequencing at Plasmidsaurus for 20 million raw 3’ end counting reads.

### Propidium iodide DNA staining for cell cycle analysis

MDA-MB-231 transduced with doxycycline-inducible IRF1 were cultured to 40% confluency in 6 cm dishes. Then, IRF1 expression was induced with 1 µg/ml doxycycline. The cells were collected by trypsinization 24h after the start of IRF1 expression. The harvested cells were washed with PBS, pelleted, and fixed in cold 70% ethanol at 4 °C for at least 30 min. The ethanol-fixed cells were washed twice with PBS and resuspended in 300 µl solution containing 5 µg/ml propidium iodide and 25 µg/ml RNase A in PBS. The mixture was incubated at 37 °C in the dark for 30 minutes and analyzed by flow cytometry using BD LSR Fortessa (BD Biosciences). Data from at least 10 000 cells was collected and analyzed using FlowJo.

### Incucyte

MDA-MB-231 cells were cultured to be at 50% confluency before harvesting to seed for time-lapse imaging in Incucyte S3 (Sartorius). 5,000 cells were seeded in a 96-well plate. Each condition was performed in four technical replicates in one experiment. 1 µg/ml doxycycline was used to induce IRF1 expression. The cells were imaged with phase-contrast every 3 hours using a 10x objective with S3/SX1 G/R Optical Module. The cells were segmented using Adherent Cell-by-Cell segmentation.

## Data and Code Availability

The raw data generated in this study is available in the supplementary material. The python code used for data processing in this study is publicly available and has been deposited in Github at https://github.com/Creixell-lab/, under Apache 2.0 license.

## Acknowledgements

We would like to thank the Proteomics, Research Instrumentation and Cell Services, and Flow Cytometry core facilities and the Laboratory Management Team at Cancer Research UK Cambridge Institute for their critical support and contribution to this project. We thank Reiner Schulte and Gabriela Grondys-Kotarba of the CIMR Flow Core Facility for cell sorting. We also thank all members of the Creixell lab for insightful discussions and sharing reagents. This research was funded by core support from Cancer Research UK (CRUK C9545/A29580) (P.C.) and UKRI grant EP/X042065/1 (M.O.).

## Author contributions

Conceptualization, M.O.; Methodology, S.P., A.B., M.O.; Software, M.J., M.O.; Investigation, S.P., A.B., L.K., M.L. M.O.; Writing—original draft, S.P. and M.O.; Writing—review & editing, S.P., P.C. and M.O. with contributions from all authors; Funding acquisition, M.O., and P.C.

## Supplementary material

**Supplementary Figure 1. Identification of HCF1 binding motifs.** (**A**) Alignment of published HCF1 binding motifs. (**B**) Correlation of peptide HCF1 binding scores measured in bacterial surface peptide display experiments using a pooled library of 347 D/EHxY peptides and their D/EHxA negative control variants. (**C**) Venn diagram showing the overlap between detected and previously published HCF1 binding motifs. (**D**) Plot showing the relationship between peptide display efficiency (MYC signal) and HCF1 binding in a bacterial surface display experiment using a pooled library of D/EHxY peptides. (**E**) Distribution of peptide enrichment scores in proteome-wide HBM peptide display experiment in D/EHxY consensus motifs that were outcompeted by HBM competitor peptide in an HCF1 coIP^11^, indicating functional HBMs, and motifs that were not outcompeted. The lines show median values, **** indicates p-value<0.0001 by Mann-Whitney U test. (**F**) Flow cytometry plots showing HCF1(Kelch)-EGFP and Alexa-647 anti-MYC staining of peptide displaying *E. coli* cells transformed with a proteome-wide library of N/QHxY peptides. (**G**) HCF1(Kelch)-GFP binding of 259 N/QHxY peptides from the human proteome measured in bacterial surface display.

**Supplementary Figure 2. Bacterial surface display experiments for deep mutational scanning of HCF1 binding motifs.** (**A**) Superimposed AlphaFold3 models of 10 D/EHxY peptides (shown in Fig 2B) bound to the Kelch domain of HCF1. (**B**) Flow cytometry plots showing the display of HCF1 binding peptides fused to eCPX-MYC on the *E. coli* surface and staining of these bacteria with Kelch domain of HCF1 fused to EGFP. A representative example of two biological replicates is shown. (**C**) Correlation between peptide binding scores (log2 fold change of peptide frequency in GFP-positive cell population from the frequency in input population) in two peptide display experiments with KANSL3, DIDO1, SETD1A and UL48 DMS libraries. (**D**) The wild-type-corrected binding scores of UL48 motif DMS peptides and their display efficiencies measured by enrichment in MYC-positive sorted cell population. (**E**) Scatter plot of measured peptide binding scores and display scores (enrichment in MYC positive population) in UL48 DMS peptides. (**F**) Binding curve of HCF1(Kelch) domain to FITC-KANSL3 HCF1 binding motif peptide in fluorescence polarisation. Data is average from two independent experiments. (**G**) Log2 fold change values of wild-type and mutant DIDO1 peptides from DIDO1 DMS peptide display experiment. (**H**) Correlation of DIDO1 wild-type and mutant peptide HCF1 binding scores measured in bacterial surface display DMS experiment and their binding affinities measured by competitive FP. (**I**) Sequence logo showing the averaged positional amino acid preferences from DMS of 4 HCF1 binding peptides in Fig. 2C.

**Supplementary Figure 3.** Improving E2F1 peptide binding with mutations in non-core positions. (**A**) Flow cytometry plots showing the peptide display (Alexa-647-MYC) and HCF1(Kelch)-GFP binding of *E. coli* cells expressing the indicated E2F1 motifs. A representative example of two biological replicates is shown. (**B**) Logos of D/EHxY peptides over- or underperforming their predicted binding scores in HCF1 binding.

**Supplementary Figure 4. Exploring the motif interactome of HCF1.** (**A**) Protein-protein interaction network analysis of 101 proteins that interact with HCF1 using the Kelch domain motif-binding pocket identified in^11^. This reveals 18 proteins that do not contain a D/EHxY motif and that are not known interactors of other proteins with HCF1 binding motifs. (**B**) AlphaFold3 modeling results of the 18 proteins identified in panel ‘A’ binding to the Kelch domain of HCF1. (**C**) Flow cytometry plots showing the display of a proteome-wide library of D/EHxxY peptides fused to eCPX-MYC on the *E. coli* surface and staining of these bacteria with Kelch domain of HCF1 fused to EGFP. A representative example of two biological replicates is shown. (**D**) Peptide enrichment log2 fold change in a display experiment using a proteome-wide library of D/EHxxY peptides and their D/EHxxA mutant variants. (**E**) Correlation of peptide HCF1 binding scores measured in bacterial surface peptide display experiments using a pooled library of D/EHxxY peptides and their D/EHxxA negative control variants.

**Supplementary Figure 5. HCF1 binding motif identity in IRF1 affects IRF1 transcriptional activity.** (**A**) Proliferation time-series of MDA-MB-231 cells expression doxycycline-inducible wild-type, Y164A (non-binding), or a high-affinity HCF1-binding IRF1. The cell numbers were monitored in Incucyte. The values are average with standard deviation error bars from 8 replicates. (**B**) The effect of overexpressing different IRF1 variants on the transcription of Interferon response genes in MDA-MB-231 cells measured by RNAseq. (**C**) RNAseq reveals differential expression of genes in MDA-MB-231 cells overexpression wild-type IRF1 and IRF1, where the native HCF1 binding motif has been replaced with a stronger motif from KANSL3. (**D**) Comparison of the change in expression of genes related to TNFα signalling via NFkB and epithelial mesenchymal transition in MDA-MB-231 overexpressing wild-type IRF1 to IRF1-KANSL3. Data is from RNAseq experiments.

**Supplementary Figure 6. Kelch domain of HCF1 as substrate recruitment adapter for OGT.** (**A**) The impact of KANSL3 DHxY competitor peptide expression on the abundance of O-GlcNAc modification in whole cell lysate. The competitor peptide was expressed for 24h in HCT116 cells.

## Supplementary Tables

**Supplementary Table S1.** Proteome-wide HCF1 binding motif identification peptide display experiments data.

**Supplementary Table S2.** HCF1 binding motif deep mutational scanning data.

**Supplementary Table S3.** IRF1 induction RNAseq data.

**Supplementary Table S4.** Proteomics data of a-OGlcNAc co-immunoprecipitation experiments.

**Supplementary Table S5.** Gene set enrichment analysis of proteins whose O-GlcNAcylation decreases with HCF1 docking interaction inhibition.

## References

1. Wilson, A. C., LaMarco, K., Peterson, M. G. & Herr, W. The VP16 accessory protein HCF is a family of polypeptides processed from a large precursor protein. Cell 74, 115–125 (1993).

2. Wu, T. J., Monokian, G., Mark, D. F. & Wobbe, C. R. Transcriptional activation by herpes simplex virus type 1 VP16 in vitro and its inhibition by oligopeptides. Mol. Cell. Biol. 14, 3484–3493 (1994).

3. Luciano, R. L. & Wilson, A. C. HCF-1 Functions as a Coactivator for the Zinc Finger Protein Krox20. J. Biol. Chem. 278, 51116–51124 (2003).

4. Wilson, A. C., Freiman, R. N., Goto, H., Nishimoto, T. & Herr, W. VP16 targets an amino-terminal domain of HCF involved in cell cycle progression. Mol. Cell. Biol. 17, 6139–6146 (1997).

5. Xiang, P. et al. HCF-1 promotes cell cycle progression by regulating the expression of CDC42. Cell Death Dis. 11, 907 (2020).

6. Tyagi, S., Chabes, A. L., Wysocka, J. & Herr, W. E2F Activation of S Phase Promoters via Association with HCF-1 and the MLL Family of Histone H3K4 Methyltransferases. Mol. Cell 27, 107–119 (2007).

7. Ruan, H.-B. et al. O-GlcNAc Transferase/Host Cell Factor C1 Complex Regulates Gluconeogenesis by Modulating PGC-1α Stability. Cell Metab. 16, 226–237 (2012).

8. Han, J. W. et al. Nuclear factor-erythroid-2 related transcription factor-1 (Nrf1) is regulated by O-GlcNAc transferase. Free Radic. Biol. Med. 110, 196–205 (2017).

9. Minocha, S. et al. Epiblast-specific loss of HCF-1 leads to failure in anterior-posterior axis specification. Dev. Biol. 418, 75–88 (2016).

10. Dejosez, M. et al. Ronin/Hcf-1 binds to a hyperconserved enhancer element and regulates genes involved in the growth of embryonic stem cells. Genes Dev. 24, 1479–1484 (2010).

11. Örd, M. et al. Uncovering cancer dependencies in peptide-interacting protein pockets. 2026.03.13.711608 Preprint at 10.64898/2026.03.13.711608 (2026).

12. Thomas, L. R. et al. Interaction of MYC with host cell factor-1 is mediated by the evolutionarily conserved Myc box IV motif. Oncogene 35, 3613–3618 (2016).

13. Wang, H., Yu, M., Yang, C. & Li, Q. Upregulation of HCFC1 expression promoted hepatocellular carcinoma progression through inhibiting cell cycle arrest and correlated with immune infiltration. J. Cancer 14, 1381–1397 (2023).

14. Srivastava, P. P. et al. Epigenetic co-regulator HCF-1 promotes lung cancer via O-GlcNAcylation-dependent pathways. Mol. Ther. Oncol. 33, 201046 (2025).

15. Kim, S. H. et al. O-linked-N-acetylglucosamine transferase is associated with metastatic spread of human papillomavirus E6 and E7 oncoproteins to the lungs of mice. Biochem. Biophys. Res. Commun. 483, 793–802 (2017).

16. Kumar, M. et al. The Eukaryotic Linear Motif resource: 2022 release. Nucleic Acids Res. 50, D497 (2022).

17. Machida, Y. J., Machida, Y., Vashisht, A. A., Wohlschlegel, J. A. & Dutta, A. The Deubiquitinating Enzyme BAP1 Regulates Cell Growth via Interaction with HCF-1. J. Biol. Chem. 284, 34179–34188 (2009).

18. Zargar, Z. & Tyagi, S. Role of host cell factor-1 in cell cycle regulation. Transcription 3, 187–192 (2012).

19. Daou, S. et al. Crosstalk between O-GlcNAcylation and proteolytic cleavage regulates the host cell factor-1 maturation pathway. Proc. Natl. Acad. Sci. U. S. A. 108, 2747–2752 (2011).

20. Rucli, S. et al. Functional genomic profiling of O-GlcNAc reveals its context-specific interplay with RNA polymerase II. Genome Biol. 26, 69 (2025).

21. Kapuria, V., Ayushma, Herr, W. & Minocha, S. HCF-1 as a key modulator of OGT function and O-GlcNAcylation in the liver. Sci. Rep. 15, 28328 (2025).

22. Rice, J. J. & Daugherty, P. S. Directed evolution of a biterminal bacterial display scaffold enhances the display of diverse peptides. Protein Eng. Des. Sel. PEDS 21, 435–442 (2008).

23. Li, A., Voleti, R., Lee, M., Gagoski, D. & Shah, N. H. High-throughput profiling of sequence recognition by tyrosine kinases and SH2 domains using bacterial peptide display. eLife 12, e82345 (2023).

24. Lee, C. Y. et al. Systematic discovery of protein interaction interfaces using AlphaFold and experimental validation. Mol. Syst. Biol. 20, 75–97 (2024).

25. Örd, M. et al. High-throughput investigation of cyclin docking interactions reveals the complexity of motif binding determinants. Nat. Commun. 16, 7622 (2025).

26. Abramson, J. et al. Accurate structure prediction of biomolecular interactions with AlphaFold 3. Nature 630, 493–500 (2024).

27. Krystkowiak, I. & Davey, N. E. SLiMSearch: a framework for proteome-wide discovery and annotation of functional modules in intrinsically disordered regions. Nucleic Acids Res. 45, W464—-W469 (2017).

28. Kano, A., Haruyama, T., Akaike, T. & Watanabe, Y. IRF-1 is an essential mediator in IFN-gamma-induced cell cycle arrest and apoptosis of primary cultured hepatocytes. Biochem. Biophys. Res. Commun. 257, 672–677 (1999).

29. Armstrong, M. J. et al. Interferon Regulatory Factor 1 (IRF-1) induces p21(WAF1/CIP1) dependent cell cycle arrest and p21(WAF1/CIP1) independent modulation of survivin in cancer cells. Cancer Lett. 319, 56–65 (2012).

30. Dror, N. et al. Identification of IRF-8 and IRF-1 target genes in activated macrophages. Mol. Immunol. 44, 338–346 (2007).

31. Song, R. et al. IRF1 governs the differential interferon-stimulated gene responses in human monocytes and macrophages by regulating chromatin accessibility. Cell Rep. 34, (2021).

32. Mody, A. C., Ramirez, D. H. & Woo, C. M. Targeted protein O-GlcNAc reveals transcriptional functions for O-GlcNAc. Cell Chem. Biol. 32, 1486–1502.e7 (2025).

33. Özcan, S., Andrali, S. S. & Cantrell, J. E. L. Modulation of transcription factor function by O-GlcNAc modification. Biochim. Biophys. Acta BBA - Gene Regul. Mech. 1799, 353–364 (2010).

34. Comer, F. I. & Hart, G. W. O-GlcNAc and the control of gene expression. Biochim. Biophys. Acta BBA - Gen. Subj. 1473, 161–171 (1999).

35. Wulff-Fuentes, E. et al. The human O-GlcNAcome database and meta-analysis. Sci. Data 8, 25 (2021).

36. Seo, H. G. et al. Identification of the nuclear localisation signal of O-GlcNAc transferase and its nuclear import regulation. Sci. Rep. 6, 34614 (2016).

37. Chang, H.-M. & Yeh, E. T. H. SUMO: From Bench to Bedside. Physiol. Rev. 100, 1599–1619 (2020).

38. Sun, L. et al. HCFC2 is needed for IRF1- and IRF2-dependent Tlr3 transcription and for survival during viral infections. J. Exp. Med. 214, 3263–3277 (2017).

39. Palopoli, N., González Foutel, N. S., Gibson, T. J. & Chemes, L. B. Short linear motif core and flanking regions modulate retinoblastoma protein binding affinity and specificity. Protein Eng. Des. Sel. PEDS 31, 69–77 (2018).

40. Subbanna, M. S., Winters, M. J., Örd, M., Davey, N. E. & Pryciak, P. M. A quantitative intracellular peptide binding assay reveals recognition determinants and context dependence of short linear motifs. J. Biol. Chem. 108225 (2025) doi:10.1016/j.jbc.2025.108225.

41. Putta, S. et al. Structural basis for tunable affinity and specificity of LxCxE-dependent protein interactions with the retinoblastoma protein family. Struct. Lond. Engl. 1993 30, 1340–1353.e3 (2022).

42. Benz, C. et al. Defining short linear motif binding determinants by phage display-based deep mutational scanning. Protein Sci. Publ. Protein Soc. 34, e70174 (2025).

43. Yang, W. H. et al. NFkappaB activation is associated with its O-GlcNAcylation state under hyperglycemic conditions. Proc. Natl. Acad. Sci. U. S. A. 105, 17345–17350 (2008).

44. Chu, C.-S. et al. O-GlcNAcylation regulates EZH2 protein stability and function. Proc. Natl. Acad. Sci. U. S. A. 111, 1355–1360 (2014).

45. Andrali, S. S., Qian, Q. & Özcan, S. Glucose Mediates the Translocation of Neurod1 by O-Linked Glycosylation. J. Biol. Chem. 282, 15589–15596 (2007).

46. Housley, M. P. et al. O-GlcNAc Regulates FoxO Activation in Response to Glucose. J. Biol. Chem. 283, 16283–16292 (2008).

47. Mannino, M. P. & Hart, G. W. The Beginner’s Guide to O-GlcNAc: From Nutrient Sensitive Pathway Regulation to Its Impact on the Immune System. Front. Immunol. 13, (2022).

48. He, X. et al. O-GlcNAcylation and stablization of SIRT7 promote pancreatic cancer progression by blocking the SIRT7-REGγ interaction. Cell Death Differ. 29, 1970–1981 (2022).

49. Le Minh, G., Esquea, E. M., Young, R. G., Huang, J. & Reginato, M. J. On a sugar high: Role of O-GlcNAcylation in cancer. J. Biol. Chem. 299, 105344 (2023).

50. Popay, T. M. et al. MYC regulates ribosome biogenesis and mitochondrial gene expression programs through its interaction with host cell factor-1. eLife 10, e60191 (2021).

51. Goddard, T. D. et al. UCSF ChimeraX: Meeting modern challenges in visualization and analysis. Protein Sci. Publ. Protein Soc. 27, 14–25 (2018).

52. Szklarczyk, D. et al. The STRING database in 2023: protein-protein association networks and functional enrichment analyses for any sequenced genome of interest. Nucleic Acids Res. 51, D638–D646 (2023).

53. Otwinowski, J., McCandlish, D. M. & Plotkin, J. B. Inferring the shape of global epistasis. Proc. Natl. Acad. Sci. 115, E7550–E7558 (2018).

