## Supplementary Figure 6 for "HCF1 orchestrates O-GlcNAcylation and affinity-dependent transcription through extended molecular determinants and register-shifted binding"

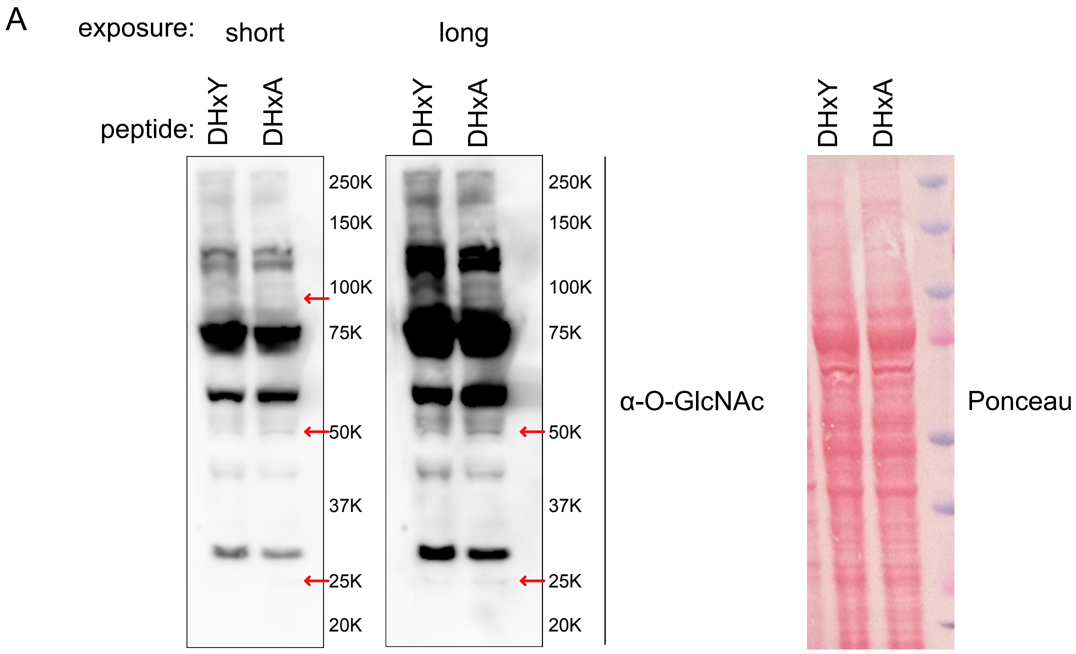


**Supplementary Figure 6. Kelch domain of HCF1 as substrate recruitment adapter for OGT.** (**A**) The impact of KANSL3 DHxY competitor peptide expression on the abundance of O-GlcNAc modification in whole cell lysate. The competitor peptide was expressed for 24h in HCT116 cells.
