## Supplementary Figure 1 for "HCF1 orchestrates O-GlcNAcylation and affinity-dependent transcription through extended molecular determinants and register-shifted binding"

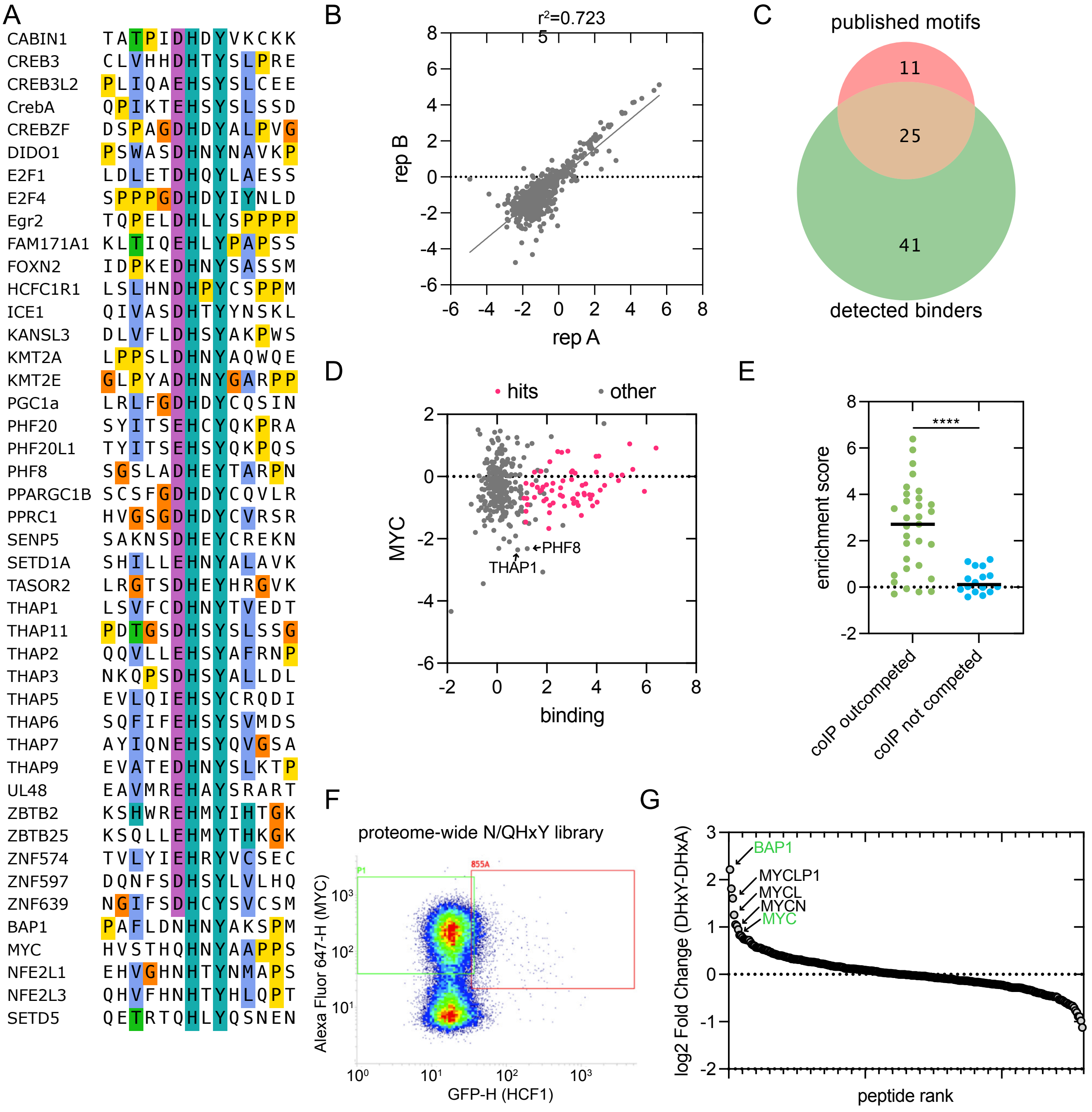


**Supplementary Figure 1. Identification of HCF1 binding motifs.** (**A**) Alignment of published HCF1 binding motifs. (**B**) Correlation of peptide HCF1 binding scores measured in bacterial surface peptide display experiments using a pooled library of 347 D/EHxY peptides and their D/EHxA negative control variants. (**C**) Venn diagram showing the overlap between detected and previously published HCF1 binding motifs. (**D**) Plot showing the relationship between peptide display efficiency (MYC signal) and HCF1 binding in a bacterial surface display experiment using a pooled library of D/EHxY peptides. (**E**) Distribution of peptide enrichment scores in proteome-wide HBM peptide display experiment in D/EHxY consensus motifs that were outcompeted by HBM competitor peptide in an HCF1 coIP11, indicating functional HBMs, and motifs that were not outcompeted. The lines show median values, **** indicates p-value<0.0001 by Mann-Whitney U test. (**F**) Flow cytometry plots showing HCF1(Kelch)-EGFP and Alexa-647 anti-MYC staining of peptide displaying E. coli cells transformed with a proteome-wide library of N/QHxY peptides. (**G**) HCF1(Kelch)-GFP binding of 259 N/QHxY peptides from the human proteome measured in bacterial surface display.
