## Supplementary Figure 2 for "HCF1 orchestrates O-GlcNAcylation and affinity-dependent transcription through extended molecular determinants and register-shifted binding"

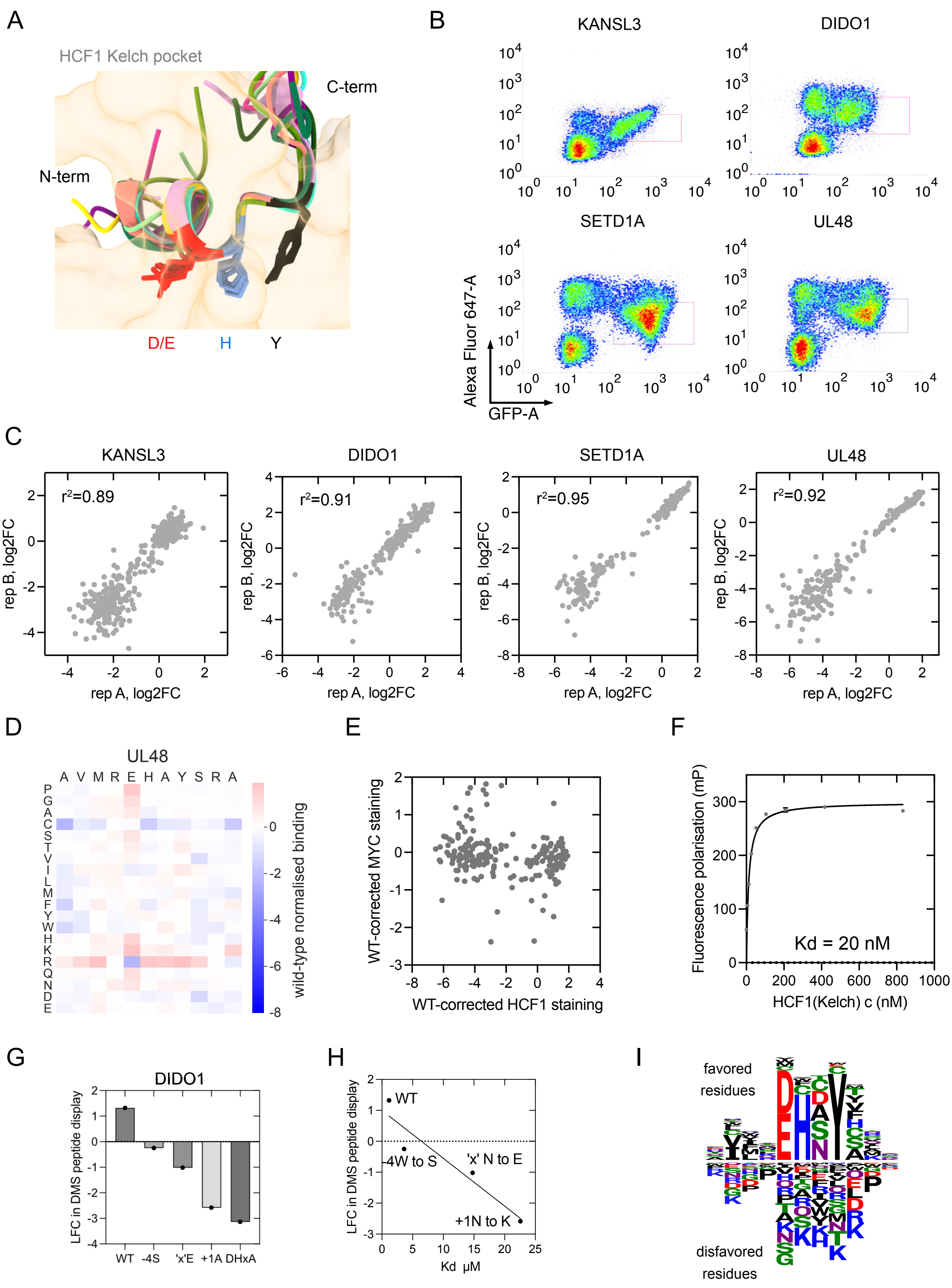


**Supplementary Figure 2. Bacterial surface display experiments for deep mutational scanning of HCF1 binding motifs.** (**A**) Superimposed AlphaFold3 models of 10 D/EHxY peptides (shown in Fig 2B) bound to the Kelch domain of HCF1. (**B**) Flow cytometry plots showing the display of HCF1 binding peptides fused to eCPX-MYC on the E. coli surface and staining of these bacteria with Kelch domain of HCF1 fused to EGFP. A representative example of two biological replicates is shown. (**C**) Correlation between peptide binding scores (log2 fold change of peptide frequency in GFP-positive cell population from the frequency in input population) in two peptide display experiments with KANSL3, DIDO1, SETD1A and UL48 DMS libraries. (**D**) The wild-type-corrected binding scores of UL48 motif DMS peptides and their display efficiencies measured by enrichment in MYC-positive sorted cell population. (**E**) Scatter plot of measured peptide binding scores and display scores (enrichment in MYC positive population) in UL48 DMS peptides. (**F**) Binding curve of HCF1(Kelch) domain to FITC-KANSL3 HCF1 binding motif peptide in fluorescence polarisation. Data is average from two independent experiments. (**G**) Log2 fold change values of wild-type and mutant DIDO1 peptides from DIDO1 DMS peptide display experiment. (**H**) Correlation of DIDO1 wild-type and mutant peptide HCF1 binding scores measured in bacterial surface display DMS experiment and their binding affinities measured by competitive FP. (**I**) Sequence logo showing the averaged positional amino acid preferences from DMS of 4 HCF1 binding peptides in Fig. 2C.
