## Supplementary Figure 3 for "HCF1 orchestrates O-GlcNAcylation and affinity-dependent transcription through extended molecular determinants and register-shifted binding"

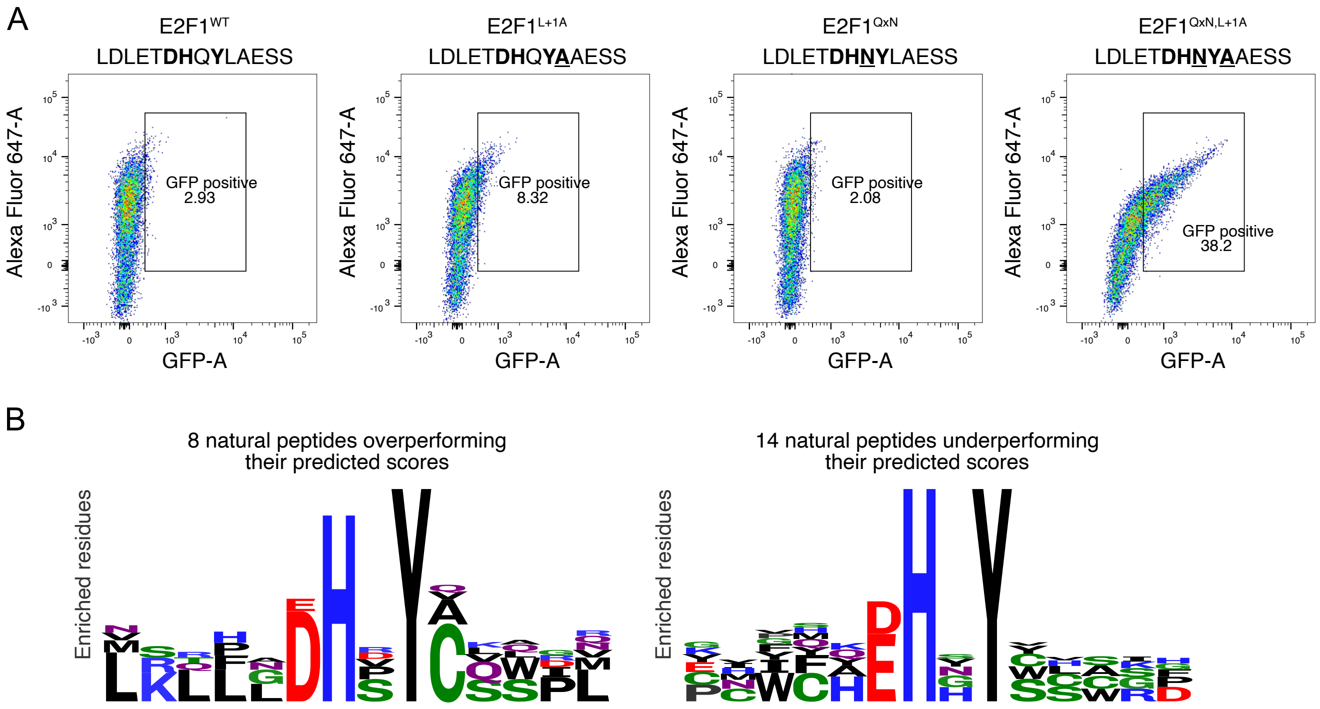


**Supplementary Figure 3. Improving E2F1 peptide binding with mutations in non-core positions.** (**A**) Flow cytometry plots showing the peptide display (Alexa-647-MYC) and HCF1(Kelch)-GFP binding of E. coli cells expressing the indicated E2F1 motifs. A representative example of two biological replicates is shown. (**B**) Logos of D/EHxY peptides over- or underperforming their predicted binding scores in HCF1 binding.
