## Supplementary Figure 4 for "HCF1 orchestrates O-GlcNAcylation and affinity-dependent transcription through extended molecular determinants and register-shifted binding"

A

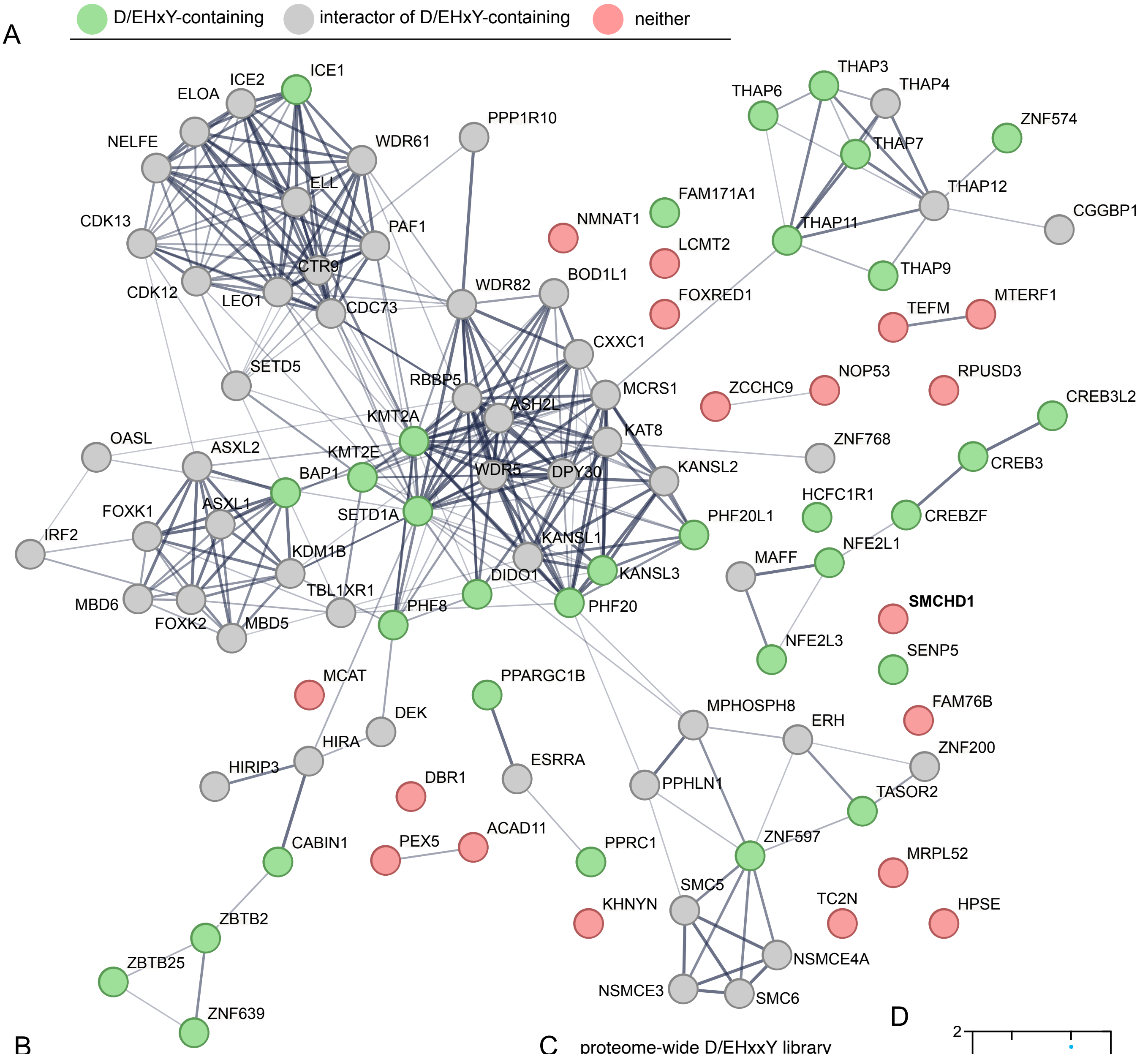


B

| Protein | in pocket? | ipTM | ipTM peptide only | peptide seq |
|---|---|---|---|---|
| SMCHD1 | yes | 0.47 | 0.89 | FES**DHSGY**VRP |
| TC2N | yes | 0.41 | 0.69 | TITLSGDER |
| ZCCHC9 | yes | 0.39 | 0.57 | WAKGMSADYE |
| TEFM | yes | 0.38 | 0.63 | ERWRC |
| NOP53 | yes | 0.28 | | ESTFQELCEG |
| MCAT | yes | 0.26 | | TLEHVDLDP |
| FOXRED1 | yes | 0.24 | | PGRSCDLLQD |
| DBR1 | yes | 0.23 | | FSDVRI |
| NMNAT1 | yes | 0.22 | | CDHQQNSPTL |
| MRPL52 | yes | 0.16 | | GPLTELPDWS |
| PEX5 | yes | 0.47 | | not IDR |
| MTERF1 | yes | 0.43 | | not IDR |
| ACAD11 | yes | 0.31 | | not IDR |
| KHNYN | yes | 0.23 | | not IDR |
| FAM76B | yes | 0.23 | | not IDR |
| RPUSD3 | yes | 0.22 | | not IDR |
| HPSE | yes | 0.14 | | not IDR |
| LCMT2 | yes | 0.14 | | not IDR |

C

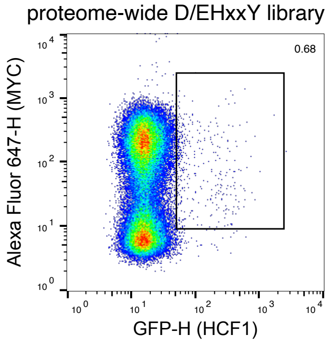


D

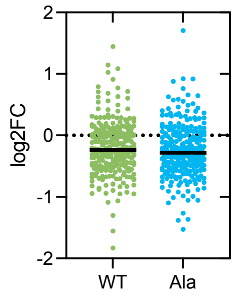


E

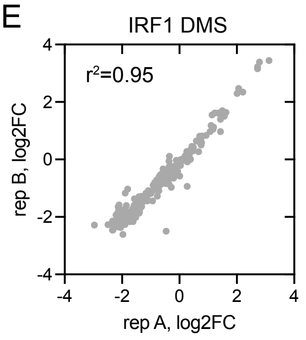


**Supplementary Figure 4. Exploring the motif interactome of HCF1.** (**A**) Protein-protein interaction network analysis of 101 proteins that interact with HCF1 using the Kelch domain motif-binding pocket identified in11. This reveals 18 proteins that do not contain a D/EHxY motif and that are not known interactors of other proteins with HCF1 binding motifs. (**B**) AlphaFold3 modeling results of the 18 proteins identified in panel 'A' binding to the Kelch domain of HCF1. (**C**) Flow cytometry plots showing the display of a proteome-wide library of D/EHxxY peptides fused to eCPX-MYC on the E. coli surface and staining of these bacteria with Kelch domain of HCF1 fused to EGFP. A representative example of two biological replicates is shown. (**D**) Peptide enrichment log2 fold change in a display experiment using a proteome-wide library of D/EHxxY peptides and their D/EHxxA mutant variants. (**E**) Correlation of peptide HCF1 binding scores measured in bacterial surface peptide display experiments using a pooled library of D/EHxxY peptides and their D/EHxxA negative control variants.
