## Supplementary Figure 6 for "HCF1 orchestrates O-GlcNAcylation and affinity-dependent transcription through extended molecular determinants and register-shifted binding"

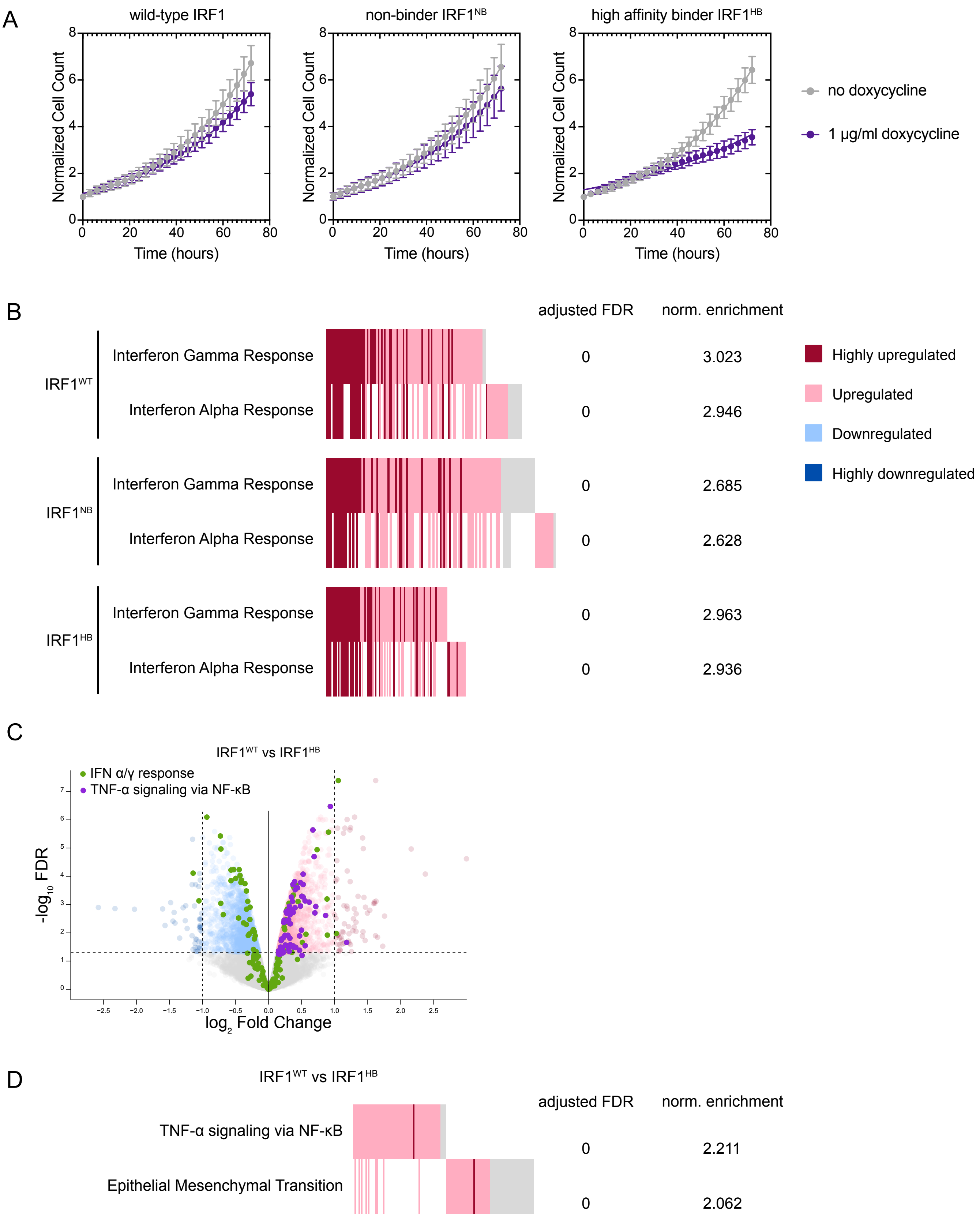


**Supplementary Figure 5. HCF1 binding motif identity in IRF1 affects IRF1 transcriptional activity.** (**A**) Proliferation time-series of MDA-MB-231 cells expression doxycycline-inducible wild-type, Y164A (non-binding), or a high-affinity HCF1-binding IRF1. The cell numbers were monitored in Incucyte. The values are average with standard deviation error bars from 8 replicates. (**B**) The effect of overexpressing different IRF1 variants on the transcription of Interferon response genes in MDA-MB-231 cells measured by RNAseq. (**C**) RNAseq reveals differential expression of genes in MDA-MB-231 cells overexpression wild-type IRF1 and IRF1, where the native HCF1 binding motif has been replaced with a stronger motif from KANSL3. (**D**) Comparison of the change in expression of genes related to TNFα signalling via NFkB and epithelial mesenchymal transition in MDA-MB-231 overexpressing wild-type IRF1 to IRF1-KANSL3. Data is from RNAseq experiments.
